# Population genomics of harbour seal *Phoca vitulina* from northern British Columbia through California and comparison to the Atlantic subspecies

**DOI:** 10.1101/2023.05.12.540438

**Authors:** Ben J. G. Sutherland, Claire Rycroft, Ashtin Duguid, Terry D. Beacham, Strahan Tucker

**Affiliations:** Sutherland Bioinformatics, Lantzville, British Columbia, Canada, V0R 2H0; Fisheries and Oceans Canada, Pacific Biological Station, 3190 Hammond Bay Road, Nanaimo, BC V9T 6N7, Canada

**Keywords:** breeding colony, genetic diversity, genotyping, harbour seal, marine mammal, *Phoca vitulina*, population genomics, RADseq, relatedness

## Abstract

The harbour seal *Phoca vitulina* is a ubiquitous pinniped species found throughout coastal waters of the Northern Hemisphere. Harbour seal impacts on ecosystem dynamics may be significant due to their high abundance and food web position. Two subspecies exist in North America, *P. v. richardii* in the Pacific Ocean, and *P. v. vitulina* in the Atlantic. Strong natal philopatry of harbour seals can result in fine-scale genetic structure and isolation-by-distance. Management of harbour seals is expected to benefit from improved resolution of seal population structure and dynamics. Here we use genotyping-by-sequencing to genotype 146 harbour seals from the eastern Pacific Ocean (i.e., British Columbia (BC), Oregon, and California) and the western Atlantic Ocean (i.e., Québec, Newfoundland, and Labrador). Using 12,742 identified variants, we confirm the recently identified elevated genetic diversity in the eastern Pacific relative to the western Atlantic and greatest differentiation between the subspecies. Further, we demonstrate that this is independent of reference genome bias or other potential technical artefacts. Coast-specific analyses with 8,933 and 3,828 variants in Pacific and Atlantic subspecies, respectively, identify divergence between BC and Oregon-California, and between Québec and Newfoundland-Labrador. Unexpected PCA outlier clusters were observed in two populations due to cryptic relatedness of individuals; subsequently, closely related samples were removed. Admixture analysis indicates an isolation-by-distance signature where Oregon seals contained some of the BC signature, whereas California did not. Additional sampling is needed in the central and north coast of BC to determine whether a discrete separation of populations exists within the region.

## Introduction

The harbour seal (*Phoca vitulina* Linnaeus 1758) is the most widely distributed pinniped species globally (Andersen & Olsen, 2010; Blanchet et al., 2021). The reported number of coastal marine harbour seal subspecies varies from three to five, but typically the recognized subspecies include *P. v. stejnegeri* (Allen 1902) in the western Pacific Ocean, *P. v. richardii* (Gray 1864) in the eastern Pacific Ocean, *P. v. vitulina* (Linnaeus 1758; formerly *P. v. concolor*; De Kay 1842) in the western Atlantic, and *P. v. vitulina* in the eastern Atlantic (Andersen & Olsen, 2010). Deep divergence exists between the Pacific and Atlantic subspecies (Liu et al., 2022), where the separation is estimated to have occurred 1.7-2.2 million years ago (Andersen & Olsen, 2010). Secondly, strong differentiation exists between western and eastern subspecies within each ocean (Liu et al., 2022; Stanley et al., 1996). Historical events and glaciations have had large influences on the present day population genetics of harbour seals (Stanley et al., 1996). A step-wise reduction in genetic diversity occurs, whereby the highest diversity is in the eastern Pacific Ocean, followed by the western Atlantic and western Pacific, and finally the lowest in the eastern Atlantic Ocean, possibly indicating colonization (Liu et al., 2022) and/ or recolonization events (Stanley et al., 1996). These historical population dynamics, along with contemporary population structure, are important to consider for species management and conservation (Olsen et al., 2014).

Harbour seals have been challenged by habitat loss, interspecific competition (Cordes et al., 2017), predation, epizootics, and human hunting or culling activity; different stressors have impacted different regions, but generally most regions have seen historical population declines, many of which have rebounded in recent years (Brown et al., 2005; DFO, 2022; Jeffries et al., 2003; Muto et al., 2021; Olesiuk, 2010). In the eastern Pacific Ocean, large-scale predator control programs and harvests in the late 1800s to mid 1900s reduced the subspecies down to approximately 10,000 seals (Olesiuk, 2010). Harbour seals were then protected in the early 1970s, after which their numbers dramatically increased to over 100,000 animals and then stabilized at the expected carrying capacity (Brown et al., 2005). By comparison, the population in eastern Canada (excluding Newfoundland) is comprised of 8,000-12,000 seals (reviewed by Blanchet et al., 2021). Throughout the range, declines and bottlenecks may have left impacts on the genetic diversity of the species (Burg et al., 1999; Westlake & O’Corry-Crowe, 2002). Severe reductions in population size in the eastern Atlantic Ocean from overhunting, environmental pollution, and epizootics, may have resulted in bottlenecks and resultant low genetic diversity (Andersen et al., 2011; Goodman, 1998; Swart et al., 1996), leading to calls for immediate monitoring in some areas (Andersen et al., 2011).

Standing genetic variation is important for rapid adaptation (Barrett & Schluter, 2008). In harbour seals, this may preserve the evolutionary potential for handling disease outbreaks or environmental perturbations (Andersen & Olsen, 2010), especially at the edges of the range where changes in climate may have a more drastic impact (Andersen et al., 2011; Blanchet et al., 2021). Genetic diversity has been directly connected to fitness in harbour seals, where lower heterozygosity was associated with increased lungworm burden in juvenile Wadden Sea harbour seals (Rijks et al., 2008). Further, birth weight and survival-to-weaning correlate with genetic diversity in harbour seal pups on Sable Island, Nova Scotia (Coltman et al., 1998). Quantifying genetic diversity of populations may help estimate adaptive potential (Barrett & Schluter, 2008), and preserving this diversity is an important conservation goal that is expected to guide management decisions.

Isolation-by-distance (IBD) is a main driver of population structure in harbour seals (Goodman, 1998; Huber et al., 2012; Liu et al., 2022; Stanley et al., 1996; Westlake & O’Corry-Crowe, 2002), with increased genetic separation occurring with discontinuous distribution of colonies (Goodman, 1998). Increased differentiation is also observed in smaller populations due to elevated impacts of genetic drift (Westlake & O’Corry-Crowe, 2002). Unexpectedly high differentiation has been observed in relatively close proximities (e.g., western and eastern Baltic Sea; Stanley et al., 1996). Harbour seals can travel great distances, but they also exhibit high philopatry to breeding colony (Goodman, 1998; Härkönen & Harding, 2001). From eastern North America to Europe, 12 regional populations have been identified, although the true number is likely greater (Andersen & Olsen, 2010), which has been further explored recently (Liu et al., 2022). Closer inspection of regions typically identifies more fine-scale structure; the United Kingdom is now delineated into northwest, northeast, and southeast populations (Olsen et al., 2017), and Ireland is delineated into northwest/north, eastern, and southwest populations (Steinmetz et al., 2023). Similarly, eastern North America was originally considered a single population, fitting with continuous clinal variation in pupping season (Temte et al., 1991), but has now been further characterized (see Liu et al., 2022). Western North America has previously been considered to be comprised of three coastal harbour seal populations, delineated as southeast Alaska through northern BC; southern BC; and Washington through California (Burg et al., 1999). A genetically distinct population was identified in Puget Sound, Washington State (Lamont et al., 1996), and inland waters of Washington State may in fact contain three discrete populations (Huber et al., 2012). Additional population structure occurs throughout Alaska, following isolation-by-distance and stepping-stone models of genetic distance (Westlake & O’Corry-Crowe, 2002). In general, most areas still require characterization of localized regions to understand population dynamics at this finer scale (Blanchet et al., 2021).

Clear identification of genetic units is essential for understanding population status and implementing management schemes (see Andersen & Olsen, 2010), which requires reliable and integrated data, possibly with demographic information and population viability analyses (Olsen et al., 2014; Olsen et al., 2017). In BC and Washington, perceived impacts of harbour seals on salmon have renewed discussions on regional population control measures (Trzcinski, 2020). Moreover, increased interest in Food, Social or Ceremonial (FSC) harvests in BC has led managers to consider regional harvest allocations, but there is incomplete information on population structure or status. In BC, harbour seals are managed as one stock (DFO, 2022). Population structure and management unit designation are particularly important to understand at a fine-scale if the population is subject to harvest (Andersen & Olsen, 2010). A coastwide assessment of population genetic structure and diversity could support the definition of units relevant for management and conservation efforts (Waples & Gaggiotti, 2006), although other considerations, such as local demography, can also be integrated into such analyses (Olsen et al., 2014). Population health is important to consider in a metapopulation context, where some healthy populations may act as sources of seals into other areas, and other populations in decline may act as sinks (Carroll et al., 2020).

Genetic studies on harbour seals have typically used allozymes (Swart et al., 1996), mitochondrial DNA (e.g., Stanley et al., 1996; Westlake & O’Corry-Crowe, 2002), and microsatellites (e.g., Goodman, 1998; Huber et al., 2012). Recently, single nucleotide polymorphisms (SNPs) by restriction site-associated DNA sequencing (RAD-seq; Baird et al., 2008; Peterson et al., 2012) were analyzed at a global scale (Liu et al., 2022). SNPs have some known benefits over microsatellites (Hauser et al., 2011), such as ease and automation of genotyping, interoperability between labs, and fewer individuals required to estimate allele frequencies (Beacham et al., 2011). SNPs also enable screening of large numbers of markers to study both neutral and adaptive variation (Andrews et al., 2016). Although high throughput methods such as RAD-sequencing require some bioinformatic expertise (Shafer et al., 2017), once variants are identified they can be integrated into amplicon panels, which can be easier to analyze to provide consistent and effective results for conservation genomics applications (Meek & Larson, 2019). Broad-scale population genetic assessments could benefit from a consistent and high throughput genotyping method, as well as systematically sampled natal sites (Huber et al., 2012).

In the present study we genotype harbour seals from broad geographic regions in the eastern Pacific Ocean (i.e., northern British Columbia (BC), Strait of Georgia (SOG), Oregon, and California), and in the western Atlantic Ocean (i.e., Newfoundland, Labrador, and eastern Quebec). We analyze population differentiation and diversity differences, shared ancestry fractions, as well as within-population relatedness, and discuss the results in relation to the recently published global analysis (Liu et al., 2022), testing both datasets for any impacts of reference bias on diversity differences. The high quality multilocus genotypes generated here are expected to be useful for the continued characterization of the population genetics and dynamics of this species.

## Methods

### Sample collection, DNA extraction, library preparation and sequencing

Harbour seal samples were obtained as skin or blood samples from collections in northern British Columbia (NBC) and the Strait of Georgia (SOG) in southern BC, Oregon (ORE), California (CAL), Québec (QUE), Newfoundland (NFL), and Labrador (LAB) (Figure 1; Table 1; Figure S1; Figure S2). Serum samples were also obtained from harbour seals in Alaska, but extractions failed to yield sufficient DNA and therefore were not included. Samples were often opportunistically collected and came from a variety of sources that included mortalities, strandings, rehabilitating captive seals, and research surveys. Agencies or organizations that collected the samples are indicated in Additional File S1, and all collections were made according to permits obtained by the individual organization as required (see details in Table S1).

**Figure 1.**
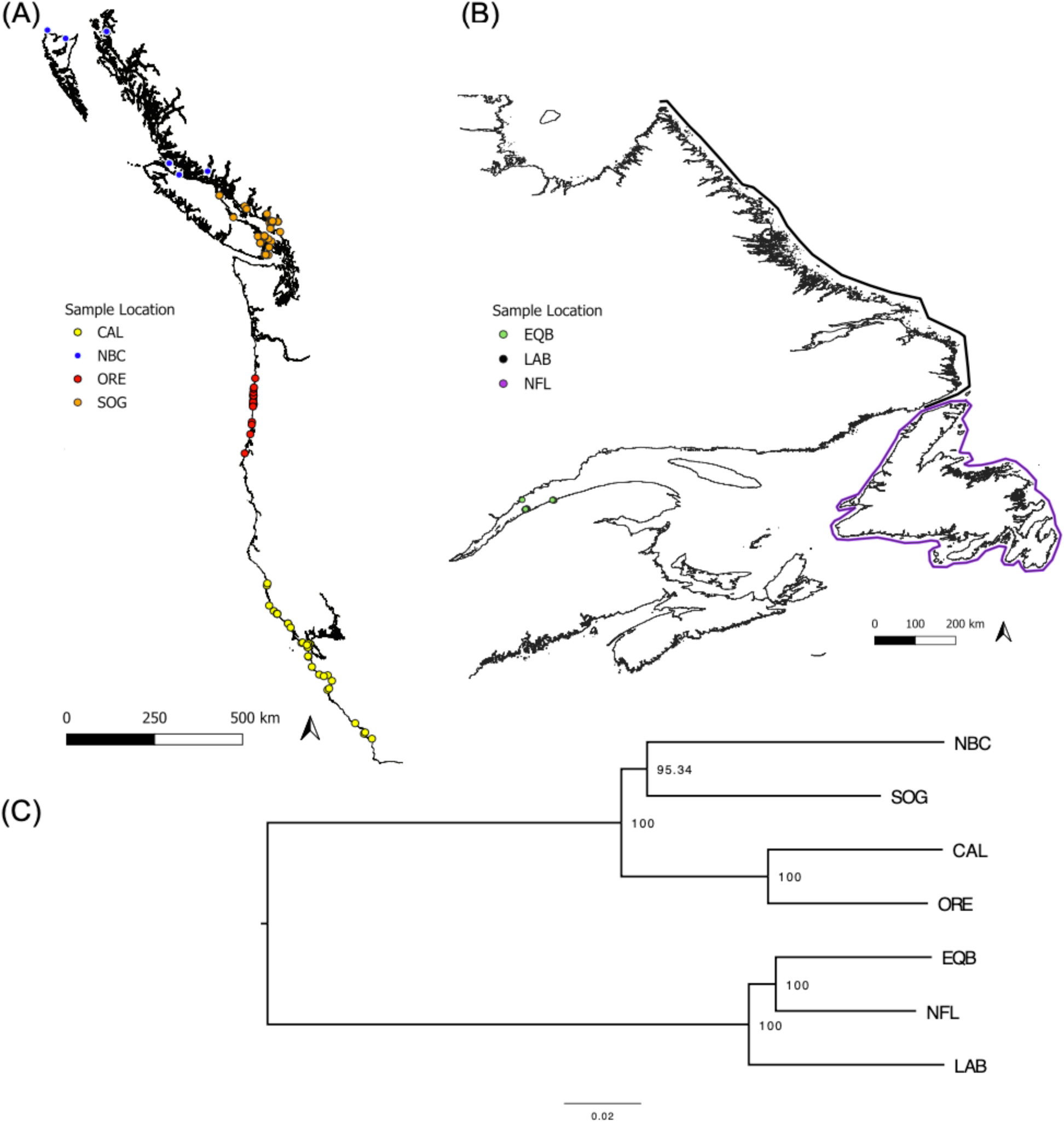
Sampling locations for the eastern Pacific (A), and western Atlantic (B) oceans, where each individual harbour seal sampled is displayed as a coloured point when individual-level sampling coordinates were available, or as a highlighted region when sample location details were not available (i.e., Labrador and Newfoundland). Genetic similarity among the populations is shown in the dendrogram (C) along with bootstrap support at the nodes. Acronyms: NBC = northern British Columbia; SOG = Strait of Georgia; CAL = California; ORE = Oregon; EQB = eastern Quebec; NFL = Newfoundland; and LAB = Labrador.

**Table 1.** Per regional collection sample size (n), total reads (M; million) and per sample average (± s.d.) per collection, total alignments (M), per sample average (± s.d.), and average alignment rate (%; ± s.d.) per collection. Values were calculated after extreme outliers (n = 2) and low coverage outliers (n = 2) were removed.

| Collection | n | Reads per collection (M) | Per sample average (M) | Alignments per collection (M) | Per sample average (M) | Alignment rate (%) |
| --- | --- | --- | --- | --- | --- | --- |
| NBC | 7 | 30.2 | $4.3 \pm 3.0$ | 27.5 | $3.9 \pm 2.8$ | $90.0 \pm 2.2$ |
| SOG | 41 | 138.6 | $3.4 \pm 1.4$ | 122.7 | $3.0 \pm 1.2$ | $88.5 \pm 1.2$ |
| ORE | 19 | 104.5 | $5.5 \pm 2.4$ | 99.5 | $5.2 \pm 2.3$ | $95.1 \pm 0.8$ |
| CAL | 30 | 108.7 | $3.6 \pm 1.9$ | 102.8 | $3.4 \pm 1.8$ | $94.1 \pm 1.8$ |
| EQB | 20 | 73.8 | $3.7 \pm 2.0$ | 63.9 | $3.2 \pm 1.8$ | $86.2 \pm 2.0$ |
| LAB | 7 | 29.0 | $4.1 \pm 2.3$ | 25.6 | $3.7 \pm 2.1$ | $87.3 \pm 3.3$ |
| NFL | 22 | 76.8 | $3.5 \pm 2.0$ | 66.5 | $3.0 \pm 1.9$ | $85.9 \pm 2.7$ |

DNA was extracted using magnetic bead extraction on a BioSprint automated DNA extraction system, following manufacturer’s instructions (QIAGEN). Extracted purified DNA was then quantified by Qubit 2.0 with the dsDNA BR assay kit (Thermo Fisher) and normalized to 20 ng/µl in 10 µl volume. The normalized DNA was submitted to the Institute of Integrative Biology and Systems (IBIS) Genomic Analyses Platform of Université Laval, where libraries were prepared using double-digest restriction-site associated DNA sequencing (ddRAD-seq). The enzymes *Nsi*I and *Msp*I were used based on *in silico* cut site assessment using SimRAD (Lepais & Weir, 2014). Libraries were sequenced on an Ion Torrent sequencer (Thermo Fisher) in batches of 20 samples per sequencing chip. Raw single-end sequence data of variable read lengths with a mode of 177 bp were obtained for downstream analyses.

### Bioinformatics

Raw sequence data were inspected for quality using fastqc (v.0.11.4; Andrews, 2010) and visualized using multiqc (v.1.14; Ewels et al., 2016). In general, the pipeline stacks_workflow (commit 868b410; E. Normandeau, GitHub) was followed, as explained below in brief. Adapters and any reads shorter than 50 bp were removed using cutadapt (v.1.9.1; Martin, 2011). Sequence data were then de-multiplexed using *process_radtags* of Stacks2 (v.2.62; Rochette et al., 2019) in parallel (Tange, 2022) trimming to a uniform length of 80 bp and de-multiplexing based on cut sites from the appropriate restriction enzymes. Samples were re-inspected for quality, then aligned to the harbour seal reference genome (i.e., GCA_004348235.1_GSC_HSeal_1.0; sourced from a female Pacific subspecies) using bwa (v.0.7.12-r1039; Li, 2013) and converted to bam format using samtools (v.1.17; Danecek et al., 2021). Numbers of reads and alignments per sample were quantified using custom scripts (see Data Availability). Two samples were found to be extreme outliers in a downstream principal components analysis (PCA) and were removed (i.e., NFL_102; NFL_121), as were two samples with fewer than 1 M reads (i.e., NBC_110, ORE_104).

Several approaches to genotyping were applied. First, the reference genome-based approach used *gstacks* with software defaults. The *populations* module was then used to filter, retaining any locus that was present in at least 70% of individuals, in all seven regional groupings, and at a minor allele frequency (MAF) ≥ 0.01. Second, to evaluate possible effects of the reference genome on genotyping of the different subspecies, *de novo* genotyping approaches were conducted within Stacks2 for each coast, using a balanced design of 18 samples from each of SOG, ORE (eastern Pacific), and EQB, and NFL (western Atlantic). The datasets were further balanced by normalizing all samples to 1.25 M reads each using seqtk’s sample function (v1.3-r106; Heng Li). The analytic pipeline for the project with all code is available on GitHub (see Data Availability, *ms_harbour_seal*).

### Population genetics

The percentage of polymorphic genomic sites, and per-population average observed heterozygosity (H_OBS_) and nucleotide diversity (π) were calculated considering all genotyped loci in Stacks2. A variant call format (VCF) file was generated using a single SNP exported per locus, and the mean genotype depth per locus per sample and per population was calculated from the VCF file using the *extract.gt* function of vcfR (Knaus & Grünwald, 2017). For downstream analysis, a genepop file was also generated including a single SNP per locus using the Stacks2 populations module, and this was read into the R environment (R Core Team, 2023) using adegenet (v.2.1.8; Jombart & Ahmed, 2011). The data were also exported from Stacks2 including all variants in radpainter format for use in fineRADstructure (Malinsky et al., 2018). The fineRADstructure analysis used software defaults and tutorials provided by the developers (see Data Availability, *ms_harbour_seal*). The single-SNP per RAD-tag dataset was converted to genlight format using dartR (v.2.0.4; Gruber et al., 2018) and coast-specific datasets were generated (i.e., Atlantic or Pacific), each of which included a coast-specific MAF filter (i.e., MAF > 0.01).

Unsupervised PCAs were generated for the global and coast-specific datasets. Due to unexpected clustering of samples within regional collections, inter-individual relatedness was estimated between samples using related (v.1.0; Pew et al., 2015) and the Ritland statistic (Ritland, 1996) to characterize potential cryptic relatedness. This analysis was conducted in the coast-wide data, as well as within PCA groupings (e.g., all BC samples) to determine the impact of population structure on the estimated relatedness. Within-population outlier pairs were identified based on the coast-wide data and defined as pairs with Ritland relatedness values above the third quartile + 1.5x the interquartile range for that population. For each outlier pair, one of the individuals was removed from the dataset until no more outlier pairs remained above the cutoff per population, as described above. After purging the putative local relatives, each dataset was filtered again for MAF (i.e., MAF > 0.01). Loci were then evaluated for Hardy-Weinberg proportions within each population using the function *hw.test* of pegas, removing any variant that was not conforming to HWP (p ≤ 0.01) in any one of the seven regional groupings. H_OBS_ and *F*_ST_ were then calculated per locus for each dataset using the *summary* function of adegenet and the function *Fst* within pegas (v.1.1; Paradis, 2010), respectively, and any locus with excess observed heterozygosity (H_OBS_ > 0.5) was removed. These datasets were then used as inputs to coast-specific PCAs, genetic differentiation assessment between populations with 95% confidence intervals of *F*_ST_ (Weir & Cockerham, 1984) in hierfstat (v.0.5-11; Goudet & Jombart, 2022), and a neighbour-joining tree using the edwards.dist distance metric (Cavalli-Sforza & Edwards, 1967) was generated using 10,000 bootstraps with the aboot function of poppr (v.2.9.3; Kamvar et al., 2014). The retained loci and individuals in the final datasets were used to subset the VCF file and vcftools (Danecek et al., 2011) was used to prepare bed, bim, and fam plink files to be used in ADMIXTURE (Alexander et al., 2009). ADMIXTURE was run using *K* values from 1-6 for the full dataset and for the east coast dataset. Optimal *K* was evaluated using a combination of lowest cross-validation error, optimal *K* metrics MedMed K, MedMean K, and MaxMed K (Puechmaille, 2016), and clumpak visualizations (Kopelman et al., 2015) as implemented in StructureSelector (Li & Liu, 2018), as well as custom R scripts (see Data Availability). Private alleles were also identified between specific groupings of samples using poppr. Analyses were conducted using functions within the GitHub repository *simple_pop_stats* with analytic R scripts available within the repository *ms_harbour_seal* (see Data Availability).

### Re-analysis of Liu et al. 2022

A recently published RAD-sequencing dataset using the *Pst*I enzyme that used genotype likelihood calls to investigate global trends in harbour seal genomic diversity (Liu et al., 2022) was downloaded using the RunSelector from the Short Read Archive (SRA) of NCBI and the SRA Toolkit (NCBI, 2023). Specifically, 14 samples from each of the following collections were downloaded: Endicott Arm (END) and Kodiak, Alaska (KOD); Île du Bic (BIC) in the St. Lawrence River, Quebec; Newfoundland (NFL); Orkney, Scotland (ORK); and Wadden Sea, Netherlands (WNL). Samples were analyzed using the reference-based genotyping methods described above, as well as by region-specific, individual *de novo* analyses. All reads were truncated to 70 bp and any read shorter than 70 bp was removed from the analysis, due to Stacks2 developers’ recommendation to keep all reads the same length for *de novo* analyses. In brief, the all-region approach retained loci present in all six populations in at least 70% of individuals per population, with global MAF ≥ 0.01. The three separate *de novo* analyses were also filtered requiring both populations in each dataset to be genotyped in at least 70% of the individuals in each population, and all retained variants to have global MAF ≥ 0.01. Population-level average H_OBS_ was calculated by the Stacks2 populations module.

## Results

### Genotyping

The 146 samples retained in the analysis had an average (± s.d.) of 3.8 ± 2.0 M reads each and were comprised in total of 561.6 M reads (see Table 1 for totals per population). No outlier samples were identified at this stage (but see Methods for earlier outlier removal). Alignment rates against the reference genome, which was generated from a Pacific subspecies female, ranged from 88.5-95.1% for Pacific samples and 85.9-87.3% for Atlantic samples. Genotyping guided by reference genome alignments used 96% of the 508.6 M primary alignments, discarding 2.8% due to low mapping quality and 1.2% due to excessive soft clipping. This resulted in the formation of 1.11 M loci (i.e., RAD-tags) comprised of 488.1 M reads with an average effective coverage of 27.5 ± 27.9x prior to filtering. Filtering for missing data and minor allele frequency (MAF; see Methods) resulted in 53,684 loci retained with a mean length of 86.6 bp. For the genotyped loci, per individual missing data was on average 2.9 ± 3.0%. These final loci were comprised of 4.6 M genomic sites, and 12,742 variants were observed, or 10,847 variants if only retaining a single SNP per locus. The mean genotype depth for all samples and loci was 35.1x, with a wide variance between samples (Figure S3), with each population ranging from a mean genotype depth of 13.3x (SOG) to 83.3x (ORE; Figure S4). For sub-species-specific analyses, data were then separated into west and east coast datasets, and once separated, each coast was filtered again for MAF, retaining 8,933 variants in the Pacific dataset and 3,828 variants in the Atlantic.

### Subspecies genetic differentiation and diversity differences

The two subspecies were found to be highly differentiated (Figure 1C), separating across PC1 (33.3% of total dataset variation; Figure 2A), similar to that recently reported (Liu et al., 2022). As an example, CAL and NFL 95% confidence interval (c.i.) *F*_ST_ was 0.40-0.42 (Table S2).

**Figure 2.**
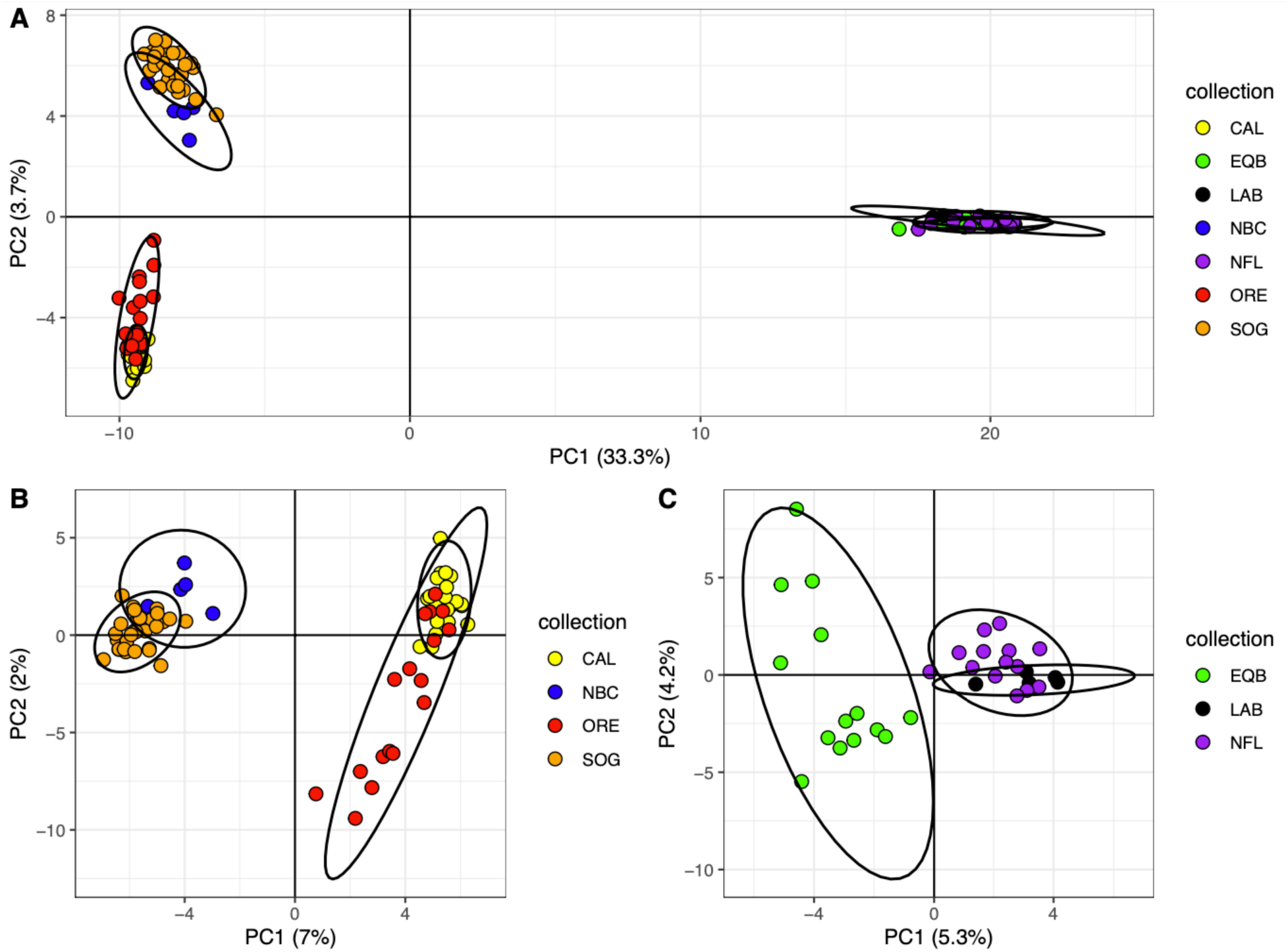
Principal components analysis following removal of putative close relatives for (A) all samples from both coasts (n = 106); (B) the Pacific subspecies (*P. v. richardii*) collections with 7,699 filtered variants; and (C) the Atlantic subspecies (*P. v. vitulina*) with 3,134 filtered variants. Ellipses represent 95% confidence level for the multivariate t-distribution.

Percentages of polymorphic genomic sites also differed greatly between the subspecies, and when considering populations with at least 15 individuals ranged from 0.180-0.194% in Pacific populations and 0.083-0.087% in Atlantic populations (see Table 2 for population-specific metrics). Observed heterozygosity (H_OBS_) was 1.9-fold higher in the Pacific than in the Atlantic subspecies for these collections (mean: 0.00045 and 0.00024, respectively), consistent with recently reported results (Liu et al., 2022).

**Table 2.** Summary statistics from genotyping 146 individuals at 12,742 variant sites in 4.6 Mbp indicates elevated percentage of polymorphic loci and observed heterozygosity (H_OBS_) in the Pacific subspecies relative to the Atlantic subspecies. This dataset is not restricted to a single SNP per RAD locus. The number of individuals with elevated genetic relatedness that were removed from each collection is shown along with the number of loci removed in the final single-SNP per locus dataset that did not conform to Hardy-Weinberg proportions. The total variants remaining in Pacific and Atlantic after all filters were 7,699 and 3,134, respectively.

| Pop | Avg. n per locus | Polymorph. variants | Percent polymorph. (%) | $H_{OBS}$ | Nucleotide diversity ($\pi$) | Related cutoff (Ritland) | Related indiv. rem. | Loci non-HWP |
| --- | --- | --- | --- | --- | --- | --- | --- | --- |
| NBC | 6.8 | 6400 | 0.138 | 0.00044 | 0.00044 | NA* | 2 | 0 |
| SOG | 39.8 | 9018 | 0.194 | 0.00043 | 0.00044 | 0.1027 | 12 | 360 |
| ORE | 18.9 | 8358 | 0.180 | 0.00046 | 0.00046 | 0.0921 | 2 | 289 |
| CAL | 29.7 | 8528 | 0.184 | 0.00045 | 0.00046 | 0.0975 | 8 | 281 |
| EQB | 19.5 | 3875 | 0.083 | 0.00024 | 0.00024 | 0.1061 | 6 | 123 |
| LAB | 6.9 | 3257 | 0.070 | 0.00025 | 0.00024 | 0.0433 | 2 | 0 |
| NFL | 21.4 | 4044 | 0.087 | 0.00024 | 0.00024 | 0.0461 | 8 | 131 |

\*no outlier pairs, applied same cutoff as SOG

To further evaluate the large differences in polymorphism between the coasts, we re-analyzed the data per coast using a consistent sample size and read depth per sample, and without the reference genome (i.e., *de novo*; see Methods). For each of the two coasts, 45 M reads were used in total, and after filters, 17,490 loci comprising 1.4 M genomic sites and 3,373 variants were observed for the Pacific subspecies (mean percent polymorphic sites = 0.183%), and 40,909 loci comprising 3.3 M genomic sites and 3,483 variants for the Atlantic subspecies (mean percent polymorphic sites = 0.085%). This demonstrates that the cause of the elevated diversity was not due to reference genome bias. Notably, in the balanced and normalized dataset, ORE showed higher observed heterozygosity than SOG (H_OBS_ = 0.00048 and 0.00039, respectively; Table S3).

A subset of the data from Liu et al. (2022) was also re-analyzed to confirm the trends viewed here. Using the reference-guided genotyping on the subset of the samples described (see Methods), Pacific H_OBS_ ranged from 0.00031 to 0.00034, whereas the western Atlantic was estimated at 0.00019 (i.e., 1.7-fold higher H_OBS_ in the Pacific), and 0.00009-0.00010 in the eastern Atlantic (Table S3). The *de novo* approach applied to a subset of the Liu et al. (2022) data resulted in a Pacific H_OBS_ of 0.00074 and Atlantic H_OBS_ of 0.00035-0.00037 (i.e., 1.8-fold higher H_OBS_ in Pacific). Collectively, this indicates that the differences in diversity reported previously and that are observed here are not due to differences in sample size, read depth, nor genetic divergence of individuals from the reference genome (i.e., reference bias).

### Cryptic relatedness within populations and genetic differentiation between populations

Coast-specific datasets were each inspected by PCA, and this showed an outlier cluster with nine individuals in the SOG, all of which were sampled in Burrard Inlet, and an outlier grouping comprised of harbour seals from the southernmost sampled region in CAL (Figure S5). Each population was then inspected for relatedness in a coastwide analysis (Figure S6), and it was determined that both PCA outlier clusters were comprised of individuals with elevated relatedness (outlier pairs Ritland metric ≥ 0.0975 and 0.1027 in CAL and SOG, respectively). Notably, when analyzing relatedness within a single regional grouping (e.g., BC only) rather than using all samples on the coast, the absolute value of relatedness metric decreased, but the same trend of outliers remained (see Figure S7).

Private alleles were identified for the Burrard Inlet cluster, which identified a private allele observed nine times in seven of the individuals (i.e., 586731_19.04), two private alleles observed six times (i.e., 729912_41.01, 807825_8.01), and three private alleles observed five times (i.e., 71641_26.04, 503353_64.04, 839341_28.01). By comparison, analyzing a haphazard selection of nine SOG individuals that were not from the Burrard Inlet cluster found no private alleles observed more than three times.

A population tree derived from the co-ancestry matrix based on shared microhaplotypes between individuals indicated two shallow branches for the Burrard Inlet cluster (Figure S8), but there was no difference between or within these two branches in the inter-individual relatedness statistic (Additional File S2), and therefore individuals within the Burrard Inlet cluster were considered approximately equally related, with no evidence for further family structure. Elevated relatedness outlier pairs were then identified and one individual from each outlier pair was removed to remove putative close relatives (see Methods; Table 2; Table S4), resulting in the retention of 73 and 33 individuals in the Pacific and Atlantic datasets, respectively (total n = 106). MAF filters were then reimposed (Pacific retained: 8,721; Atlantic retained: 3,658), and loci not conforming to HWP or with excess H_OBS_ were removed (Table 2), resulting in 7,699 and 3,134 variants retained in the Pacific and Atlantic datasets, respectively. Of these remaining variants, 56.3% and 46.5% were between 0.01-0.10 MAF in the Pacific and Atlantic harbour seal datasets, respectively (Figure 3).

**Figure 3.**
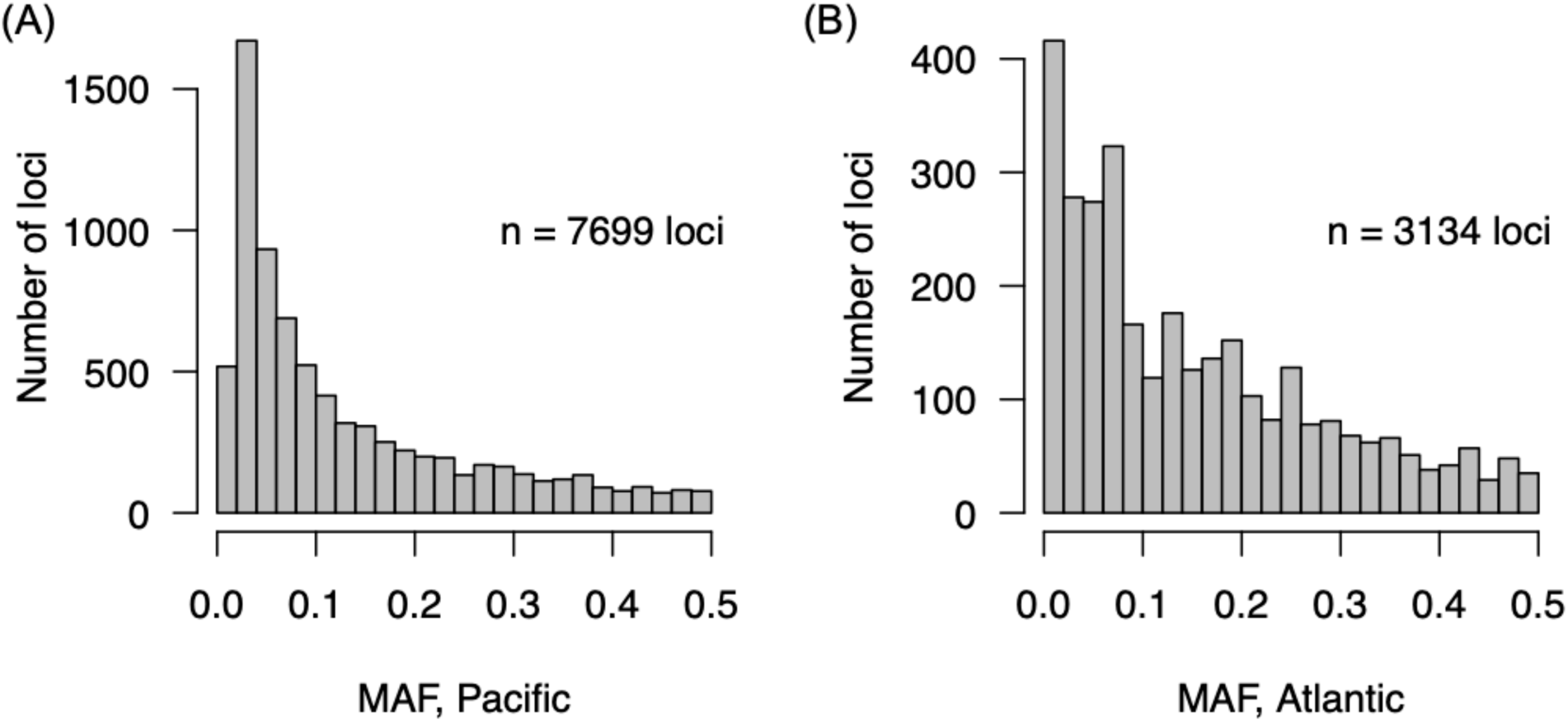
Minor allele frequency distribution for (A) Pacific or (B) Atlantic datasets after removing low MAF markers within each coast-specific dataset (i.e., retain MAF ≥ 0.01). Markers with MAF between 0.01 to 0.10 comprise 56.3% and 46.5% of the Pacific and Atlantic datasets, respectively.

The Pacific dataset shows clear differentiation between the southern and northern samples, explaining 7% of the variation across PC1 and separating SOG/NBC from ORE/CAL (Figure 2B; Figure S9 for Scree plots), with significant genetic differentiation (e.g., SOG-CAL 95% c.i. *F*_ST_ = 0.065-0.071; Table 3A). Within BC, overlap of confidence interval ellipses is observed for NBC and SOG, as well as for ORE and CAL in the USA. These collections show similar levels of differentiation, with NBC-SOG *F*_ST_ = 0.014-0.021 and ORE-CAL *F*_ST_ = 0.006-0.009 (Table 3A). Due to the low sample size of the NBC collection, the SOG-NBC *F*_ST_ value should be taken with caution. Notably, the closest SOG samples in the PCA to the NBC samples were some of the most northern samplings within the SOG collection, including Nelson Island on the Sunshine Coast (SOG_118), and Anvil Island in Howe Sound (SOG_123). Sample NBC_111, sampled at northern Haida Gwaii was one of the most distant samples geographically, and the most distant sample along PC1 from the SOG samples (Figure 2B; Additional File S3). Putative relatives (removed in final analysis), NBC_112 and NBC_113, collected from Prince Rupert and Massett (Haida Gwaii), respectively, were also grouped together with NBC_111 distantly from the SOG, but this is also expected due to the estimated relatedness to NBC_111. However, while some geographic trend is therefore apparent, given the low number of characterized samples from northern BC, there is no clear delineation yet observed in the data between the SOG and the more northern samples (Figure 2B).

**Table 3.** Genetic differentiation (Weir-Cockerham *F*_ST_) between populations in the (A) Pacific subspecies dataset comprised of 7,699 variants; and in the (B) Atlantic subspecies dataset comprised of 3,134 variants. Values in the upper and lower sections are upper and lower 95% confidence limits of the estimated *F*_ST_, respectively. Any negative values should be considered *F*_ST_ = 0, or no differentiation.

|  | NBC | SOG | ORE | CAL |
| --- | --- | --- | --- | --- |
| NBC | - | 0.021 | 0.052 | 0.064 |
| SOG | 0.014 | - | 0.057 | 0.071 |
| ORE | 0.043 | 0.052 | - | 0.009 |
| CAL | 0.055 | 0.065 | 0.006 | - |

(B)
|  | EQB | NFL | LAB |
| --- | --- | --- | --- |
| EQB | - | 0.023 | 0.032 |
| NFL | 0.016 | - | 0.002 |
| LAB | 0.020 | -0.007 | - |

The Atlantic subspecies analysis has a generally strong separation between the EQB and the LAB/NFL collections along PC1 (5.3% of the variation; Figure 2C; Figure S9). After removing putative siblings from the EQB collections, the three samples that were highest on PC2 were sampled at or near the Métis harbour seal colony (EQB), and all those lower on PC2 were from the Le Bic colony (EQB), which appears to be explaining the separation of these samples across both PC2 and PC1. The LAB and NFL collections were completely overlapping with each other and have an *F*_ST_ value overlapping zero, suggesting that these are not separate populations. The separation of EQB and NFL is however reflected in the significant *F*_ST_ between EQB and NFL (95% c.i. *F*_ST_ = 0.016-0.023; Table 3B).

Admixture analysis using data from both coasts after removing putative relatives was conducted at *K* = 1-6 (all major modes shown in Figure S10). The lowest cross-validation (cv) error occurred at *K* = 3 (median cv error = 0.3019; Figure S11), and although *K* = 4 had a slightly higher value (cv error = 0.3208), the minor modes of *K* = 4 (4/10 runs) clearly separated EQB from NFL/LAB (Figure 4; Figure S12). The major modes of *K* = 4 (6/10 runs) separated the SOG into mixed ancestry individuals. When analyzing the east coast dataset by itself, although the lowest CV error was at *K* = 1 (Figure S13), StructureSelector found MedMed *K*, MedMean *K*, and MaxMean *K* all supporting the optimal *K* = 2 in the east coast data (Figure S14), which separated EQB from LAB/NFL as per the minor modes of *K* = 4 in the both coast dataset (Figure S15). Notably, the minor modes of *K* = 4 in the both coast data indicated that the SOG samples were nearly completely comprised of the SOG ancestry, whereas the NBC samples had a small fraction of ancestry shared with the other coastal samples from ORE and CAL. The lack of ORE/CAL ancestry in the SOG was common in both the major and minor modes of *K* = 4. The ORE samples showed a small fraction of the BC (SOG) signature, whereas the CAL samples were comprised completely of the CAL ancestry, suggesting that ORE is more similar to BC than CAL is to BC. Both the PCA and the admixture analysis indicate that ORE is most similar to CAL, even though its distance from BC and CAL are approximately similar (Figure 1A).

**Figure 4.**
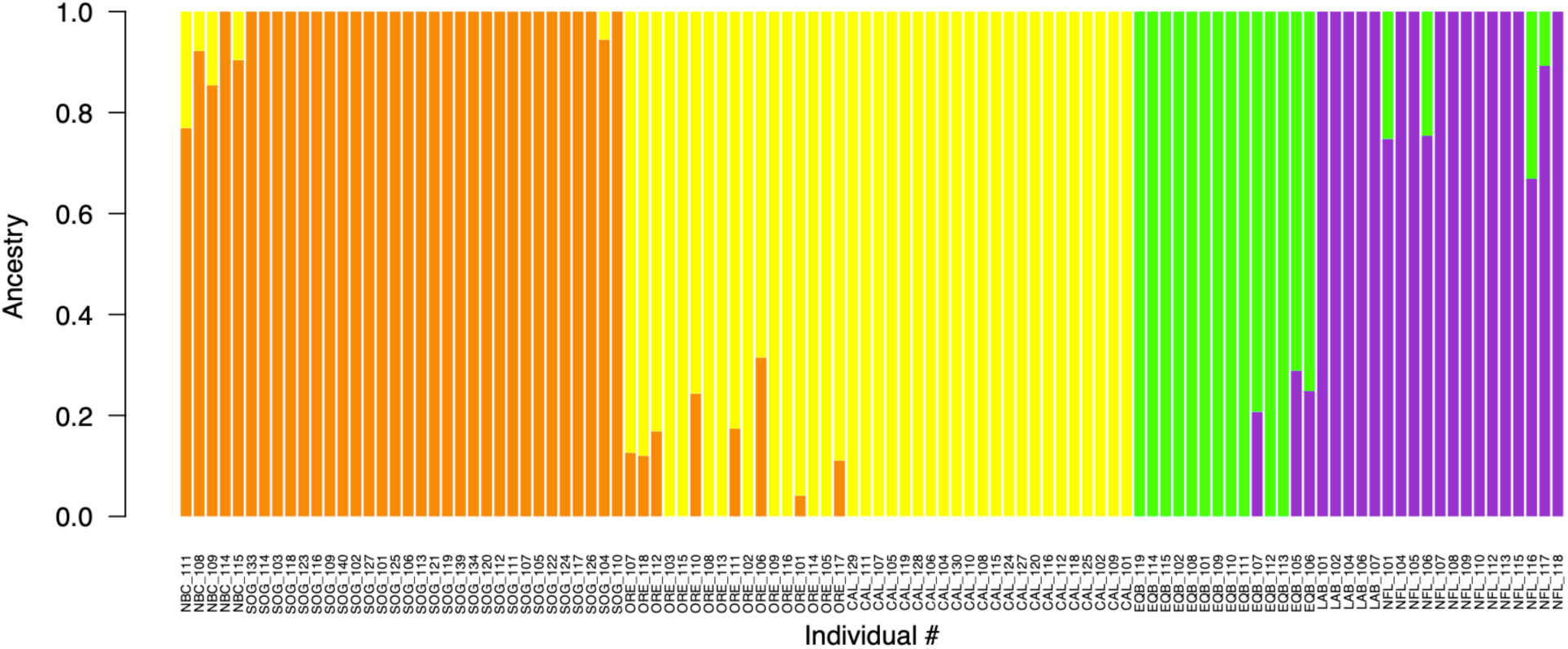
Admixture analysis with *K* = 4 including individuals from both coasts (n = 106 individuals) after removal of individuals with elevated estimated genetic relatedness. This plot represents a single run, and plots with other values of *K* and multiple runs are shown in the supplemental materials. Pacific samples (i.e., NBC, SOG, ORE, CAL) are arranged by latitude, with the leftmost samples being those that were sampled from the most northern locations. Atlantic samples are grouped only by region, as most did not have exact sampling coordinates available.

### Future work and utility of identified markers

Markers were identified here that may be useful for further characterization of population structure (i.e., loci with elevated *F*_ST_ between southern and northern BC or between BC and Oregon) or for individual identification (i.e., elevated per locus H_OBS_ in both the BC and US collections; Additional File S4). Contrasting per locus H_OBS_ between the BC and US datasets indicates high correlation in H_OBS_ between the datasets (Figure S16). These markers could be implemented to design genotyping panels as needed, and those with consistently elevated H_OBS_ in multiple populations may have utility on a coastwide scale, at least in the regions assessed here.

## Discussion

### Subspecies divergence and differential genetic diversity

Species conservation and management of harbour seal will likely depend on effective characterization of population structure and contemporary standing genetic variation (including functional variation), that is considered in the context of large-scale historical demographic events (as per Beichman et al., 2023), as well as contemporary and local demography (Olsen et al., 2014). In this study, we confirm the existence of elevated genetic diversity in the eastern Pacific subspecies of harbour seal *P. v. richardii* over the western Atlantic subspecies *P. v. vitulina* as was recently reported by Liu and co-workers (2022). We observe the elevated diversity of the Pacific subspecies in new regions including British Columbia (BC) and Oregon, whereas the previous analysis included Alaska and California for the eastern Pacific subspecies (Liu et al., 2022). Further, we evaluate and find that this trend is not caused by reference genome bias in genotyping, where individuals with increased divergence from the reference genome may exhibit biased population genetic metrics (Bohling, 2020). Reference bias is typically considered in the context of divergent species, but as more local genomes are produced, these issues may become more relevant within species (Beichman et al., 2023; Thorburn et al., 2023). Further, given the high divergence between the subspecies here, it was important to test whether *P. v. vitulina* metrics were impacted by the alignment to the *P. v. richardii* genome. The diversity difference of around 1.8x higher (evaluated by H_OBS_) in the Pacific over the Atlantic North American subspecies remains when genotyping without a reference genome (i.e., *de novo*) with a normalized number of samples and reads.

The ultimate cause of the diversity difference was proposed by Liu and co-workers (2022) to be due to stepwise founder effects during colonization from the Pacific Ocean to the western Atlantic Ocean, and then from the western Atlantic Ocean to the eastern Atlantic Ocean, with a decrease in genetic diversity at each step. Proposed west to east colonization within each basin is supported by previous work (Stanley et al., 1996), although some evidence also exists for an east to west radiation in the Pacific (Westlake & O’Corry-Crowe, 2002). Stanley and co-workers (1996) proposed that the European population may have been eliminated during the last Ice Age (∼18,000 y.a.), and that it was recolonized following the recession of the ice sheets. These estimates and additional demographic history of these populations will be important to explore with next-generation sequencing data (Beichman et al., 2023), albeit with caution given the susceptibility of demographic inference to artefacts from pipeline parameters (Shafer et al., 2017). The use of these multiple, independent RAD-seq datasets may aid in this analysis. The relatively deep sequencing and therefore highly reliable genotypes expected in the present analysis, and the numerous RAD-tags generated through restriction enzyme selection (Lepais & Weir, 2014), may enable the analysis of the allele frequency spectra of the two subspecies to characterize fluctuations in effective population size (Ne) and therefore approximate the timing of the reduced genetic diversity event (e.g., Beichman et al., 2023; Liu & Fu, 2020). Uncertainty levels around the demographic inference approaches may not be robust enough for conservation and management decisions, but results of these approaches may increase general understanding of putative and broad biogeographic hypotheses (Shafer et al., 2017).

The striking diversity differences between the subspecies may provide a model to study the effect of environmental changes and adaptive potential in the two subspecies, and their responses to environmental challenges. Furthermore, the large divergence between the Pacific and the Atlantic subspecies raises the question of whether the subspecies are in fact separate species (Liu et al., 2022). *F*_ST_ values observed by Liu et al. (2022) ranged from 0.38 to 0.69, and the authors noted that this is higher than most other marine megafaunal subspecies to date. Our results agree with these levels, and although we did not have samples from the eastern Atlantic Ocean, here the 95% lower and upper *F*_ST_ values for all contrasts between the west and east coast of North America (excluding the low sample size NBC) ranged from *F*_ST_ = 0.36 to 0.43.

### Population structure in the eastern Pacific

Harbour seals from the eastern Pacific Ocean along North America have been considered to be comprised of three coastal populations: (1) southeast Alaska through northern BC; (2) southern BC; and (3) Washington through California (Burg et al., 1999). In the present study, the separation between BC and Oregon/California is clear. Additionally, here, based on PCA, admixture analysis, and *F*_ST_ some separation is viewed among samples from southern BC and northern BC, and among samples from Oregon and California, but there are no clear break points or strong separations observed between the putative populations aside from the separation between BC and Oregon/California. Continuous distributions of breeding animals along coastlines has previously been described to have little local substructuring, although a continued increase in differentiation occurs with distance (Goodman, 1998). In the case of clinal isolation-by-distance, sampling at extreme ends of a geographical region may result in apparent clusters, whereas if the sampling was continuous throughout the range, a cline would be observed (Patterson et al., 2006). In this regard, continuous sampling along coastlines would help refine whether breakpoints exist or if a continuous distribution explains the coastal population structure. In addition to the continuous distribution of breeding animals occurring along the coast, the detection of substructure may also be impeded due to high population abundance (Burg et al., 1999). The location of a putative dividing line between southern and northern BC has yet to be identified (Burg et al., 1999). In the present study we also do not identify this line, which may be due to having too few northern BC samples genotyped here, and the location may be only identifiable with a larger survey of coastal breeding colonies. Furthermore, the samples used in the present study were mainly opportunistically collected, and this type of sampling has been proposed to blur the lines between populations as opposed to careful sampling of unweaned pups to ensure exact natal locations (Huber et al., 2012). In any case, markers used here did show some spread of samples in the PCA and some differences in admixture fractions, between northern and southern BC, and between Oregon and California. A subset of markers identified here, applied to a higher number of individuals in future sample collections using amplicon sequencing approaches for example, may help resolve some of these questions. A focus on systematic sampling of natal sites within the Central Coast region may be key in identifying a potential dividing line within BC (Burg et al., 1999), if it exists.

The Strait of Georgia (SOG) has the highest density of harbour seals on the BC coast with nearly 40,000 animals (i.e., 37% of BC population), and is used for population surveys since the implementation of protections (DFO, 2022; Majewski & Ellis, 2022). Notably, of the 12 harbour seals sampled in the Burrard Inlet area, nine were found to have high estimated genetic relatedness. These harbour seals were all pups, and were all sampled in the same summer. As female harbour seals only produce a single pup per year (Schaeff et al., 1999), it is possible that the samples could be half-sibs, sharing a sire. However, harbour seals are not expected to mate in harems, but rather in-water in a lek-type mating system (Boness et al., 2006), which makes the observation of so many putatively related pups surprising. Colonies having different levels of overall relatedness have been previously reported (Olsen et al., 2014). Regardless, the unexpected relatedness (i.e., cryptic relatedness) had significant effects on the initial PCA (Yao & Ochoa, 2023), highlighting the importance of removing putative relatives to avoid erroneous identification of population clusters (Patterson et al., 2006), given their overrepresentation of certain alleles (Elhaik, 2022). This feature of population characterization, which is expected in other methods and not only PCA (Patterson et al., 2006), should be considered in future sampling designs. It is interesting to note that the coastal BC samples in northern BC showed a fraction of admixture signature that was shared with ORE/CAL, whereas the SOG samples largely did not contain any of this signal (except one sample with a small fraction). This further suggests isolation of the SOG samples, along with the increased relatedness signatures observed. The presence or extent of any genetic separation existing between this colony (or colonies) and others is unknown, and would need follow-up sampling to understand further. The BC signature was viewed in a minor fraction of the ORE samples, but not the CAL samples, providing evidence of isolation-by-distance in harbour seals that are furthest south from BC. In harbour seals, isolation-by-distance may not explain all genetic separation, and significant differentiation can occur within only a few hundred kilometers, potentially being driven by other demographic processes (Goodman, 1998; Stanley et al., 1996).

Inland populations in Washington State that are genetically distinct from the coastal population were observed to have the most diverse mitochondrial haplotypes (Lamont et al., 1996). These populations have different pupping times and cranial morphometrics, although are geographically close, and may be more closely related to harbour seals from northern populations, including the SOG (reviewed by Lamont et al., 1996). Late birthing of pups has been observed in Puget Sound and Vancouver Island that is discontinuous with clinal variation observed further south, again suggesting some unique characteristics in this area (Temte et al., 1991). Furthermore, Burg and co-workers (1999) identified a unique mitochondrial haplotype signature in harbour seals sampled from southern Vancouver Island, but did not see this group as a separate population in their microsatellite analysis. Similarly, in the present work, we noted high frequency private alleles in the Burrard cluster of putatively related individuals prior to their removal. Unique signatures were suggested to potentially originate from a separate colonization event for these populations (Burg et al., 1999; Lamont et al., 1996). Whether the Burrard cluster is simply an artefact of chance sampling of individuals with high relatedness from a fairly genetically homogenous colony in the inlet, skewing allele frequencies and enriching rare alleles (Elhaik, 2022), or in fact is evidence of impeded gene flow, requires further study with systematically obtained samples over time. Additional samples from the inland populations of Washington State (Lamont et al., 1996) to compare with the Burrard cluster would also be important to include in future analyses. A southern California cluster was also observed, and similarly this group was found to have high estimated relatedness, and therefore was required to have close relatives removed to not appear as artefactual population structure in the PCA (Patterson et al., 2006). This provides additional support for the hypothesis that the family effect is driving the intra-collection clusters. However, it is not clear why related individuals were specifically sampled in this area through the opportunistic sampling methods applied. A caution regarding the removal of the putative close relatives is that we are unable to determine whether this is an isolated colony (without significant gene flow to neighbouring colonies); the highly related nature of the animals may simply be a characteristic of an isolated population (Waples & Anderson, 2017), and the lack of the signature of ORE or CAL in the SOG samples in the admixture analysis further indicates the isolation of this population, even in contrast with other samples further north in BC.

California harbour seals are managed separately from Oregon and coastal Washington State, but this is due to jurisdictional or political separation rather than due to genetic information (Brown et al., 2005). In the present results, the level of genetic differentiation between ORE and CAL (i.e., *F*_ST_ = 0.008-0.011) was lower than that between NBC and SOG (i.e., *F*_ST_ = 0.016-0.021), or EQB and NFL (i.e., *F*_ST_ = 0.019-0.024), but see above regarding caution for NBC allele frequencies. Given the expected gene flow between Oregon and neighbouring states, this may explain how the reduction to approximately 500 harbour seals in Oregon before protections were implemented in the early 1970s (reviewed by Brown et al., 2005) did not result in a suppression of genetic diversity in the contemporary population. Similarly, in sea otter *Enhydra lutris*, simulations suggest that accumulated genetic load through bottlenecks and low Ne decreases more rapidly with restoration of gene flow (Beichman et al., 2023). As described above, the demographic history and Ne fluctuations of Pacific and Atlantic subspecies, including within each basin, would be a valuable approach to understand how recorded historic events compare with the genetic signatures observed.

### Marker selection for an amplicon panel

Marker panels, such as a low-density 500 marker panel, are useful tools for conservation genetics (Meek & Larson, 2019), including for characterizing population structure, conducting genetic assignments and parentage, as well as for providing some estimates of genetic diversity. With the present data, markers identified here could be used to generate such a panel. To select highly differentiating markers that will separate populations along the range of the Pacific harbour seal, markers could be selected based on high *F*_ST_ both between Northern BC and SOG, and between SOG and Oregon/California. However, a limitation of this approach is that regions not sampled in the present study may be missed in such a design. Additional goals of individual identification could be considered as well, and other markers with elevated heterozygosity would be useful in this regard to include in a panel (e.g., markers with elevated H_OBS_ in both the US and Canadian populations). Removal of the highly related outlier cluster is likely to be a good approach for identifying markers, as was done here, but it is also likely valuable to include several of the high frequency private alleles identified in the Burrard cluster in the panel for broader investigation. These genetic efforts are expected to be useful for future planning of management units for the Pacific harbour seal.

## Conclusions

The present study characterizes 12,742 variants in 146 harbour seals sampled from seven different regions in the east and west coast of North America. Top differentiating or high heterozygosity variants can be used for future studies, for example to design an amplicon panel, to further characterize population structure, for parentage analysis, or to enable repeat detections of individuals. This study confirms the recent observation of elevated diversity in Pacific harbour seals over the western Atlantic subspecies, and verifies that this trend is not due to analytic artefacts such as reference bias, sequencing depth, or sample size. Although northern BC samples share some ancestry with coastal seals from Oregon and California, those sampled in the Strait of Georgia largely did not. The Oregon seals shared some of the BC ancestry signature, whereas further south, the California seals did not; this suggests isolation-by-distance, known to be a factor in population structure of harbour seals. Cryptic relatedness was observed in seals sampled in Burrard Inlet and southern California, and was subsequently removed from the dataset. Cryptic relatedness may have broader implications for future sampling efforts from colonies to characterize discrete populations. Additional samples, systematically characterized over time will be helpful to determine the extent of gene flow between putative colonies. Samples of expected parents could also help to resolve the nature of the relationships among samples in colonies through parentage analyses. Future studies using the multiple independent datasets now available could explore the demographic history of both subspecies to determine timing of Ne fluctuations on either coast to further investigate the causes of the striking differences in genomic diversity between the subspecies.

## Supporting information

Supplemental Results

## Acknowledgements

This work was supported by Fisheries and Oceans Canada Pinniped Research Program and the Species At Risk Program. The authors are grateful to sample providers, including Garry Stenson of DFO Atlantic (Newfoundland and Labrador), Mike Hammill and Xavier Bordeleau of DFO Québec (eastern Québec), the Marine Mammal Rescue Centre, Vancouver and DFO Pacific (Strait of Georgia, northern BC), James Rice of the Oregon Marine Mammal Stranding Network (Oregon), Barbie Halaska of the Marine Mammal Center of Sausalito (California), and Lauri Jemison and Lori Polasek from Alaska Department of Fish and Game (ADFG). Thanks to the Molecular Genetics Lab of DFO Pacific for technical support and discussions. Thanks to Jamieson Gorrell for discussions on admixture analyses. Thanks to Thierry Gosselin, Morten Tange Olsen, four anonymous reviewers, and the subject Editor for their constructive comments on earlier versions of the manuscript.

## Data Availability

All code required to analyze the project: https://github.com/bensutherland/ms_harbour_seal Functions used for analysis: https://github.com/bensutherland/simple_pop_stats Demultiplexed sequence data is available on NCBI through the SRA within BioProject PRJNA948428, BioSamples SAMN33902654-SAMN33902801.

Associated files on FigShare including compressed GitHub analysis pipeline, sample information file needed for *stacks_workflow*, genotypes for all populations from the reference-guided analysis (all samples or normalized) in genepop and VCF formats (both are single SNP per locus) as well as RADpainter input (microhaplotypes), and the balanced, normalized, *de novo* genotyping genepops (single SNP per locus) and VCF files (multiple SNP per locus): https://doi.org/10.6084/m9.figshare.22290472

## Benefit-sharing statement

This research addresses the population genetics of harbour seals in the eastern Pacific and western Atlantic. We have ensured that all raw data, output genotype data, and metadata is provided, as well as well-documented scripts and instructions for data analyses. All individuals and/or organizations who provided samples for the project are included in the Acknowledgements section.

## Supplemental Information

In addition to the Additional Files shown below, the Supplemental Results section contains supplemental figures and tables.

Additional File S1. All sample metadata including location, sex, and age information.

Additional File S2. Pairwise relatedness statistics displaying histogram of all pairwise values and the top interindividual relatedness pairs for both Pacific and Atlantic datasets.

Additional File S3. Coast-specific analyses PCA scores, sample IDs, location, and GPS coordinates. Additional File S4. Per locus marker statistics for (a) Pacific populations and (b) Atlantic populations, both with all putative relatives removed, as well as between (c) SOG and NBC and (d) SOG and ORE.

