## Supplemental Results for "Population genomics of harbour seal *Phoca vitulina* from northern British Columbia through California and comparison to the Atlantic subspecies"

### Table of Contents:


| | |
|---|---|
| **Figure S1. Sample map, BC** | Page 2 |
| **Figure S2. Sample map, Quebec** | Page 3 |
| **Figure S3. Genotype depth, indiv.** | Page 4 |
| **Figure S4. Genotype depth, collection** | Page 5 |
| **Figure S5. Coast-specific PCA, all indiv.** | Page 6 |
| **Figure S6. Relatedness between pairs** | Page 7 |
| **Figure S7. Relatedness, BC-only** | Page 8 |
| **Figure S8. Shared microhaplotypes** | Page 9 |
| **Figure S9. Scree plots for coast-specific PCA** | Page 10 |
| **Figure S10. Admixture proport., both coasts** | Page 11 |
| **Figure S11. Admixture CV error, both coasts** | Page 12 |
| **Figure S12. Admixture $K$ = 4, both coasts** | Page 13 |
| **Figure S13. Admixture CV error, east coast** | Page 14 |
| **Figure S14. Optimal $K$ selection, east coast** | Page 15 |
| **Figure S15. Admixture proport., east coast** | Page 16 |
| **Figure S16. Per locus $F_{ST}$ and $H_{OBS}$** | Page 17 |
| **Table S1. Collection and permit details** | Page 18 |
| **Table S2. Mean $F_{ST}$ between all collections** | Page 19 |
| **Table S3. Reference bias summary statistics** | Page 20 |
| **Table S4. Removed indiv. (relatedness)** | Page 21 |

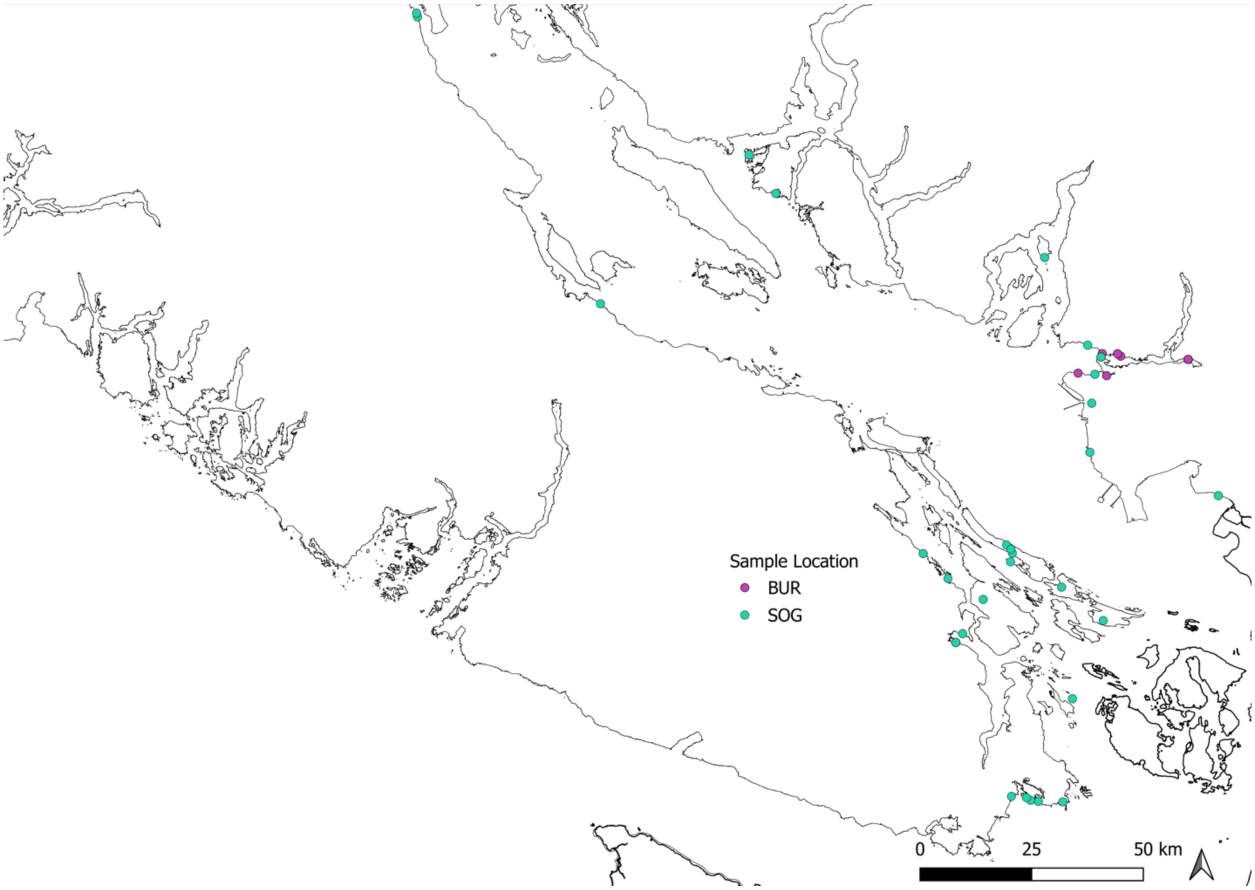


**Figure S1.** Detailed sample location map for southern BC, highlighting sampling locations for the Burrard Inlet cluster (pink) and other southern BC collections (green).

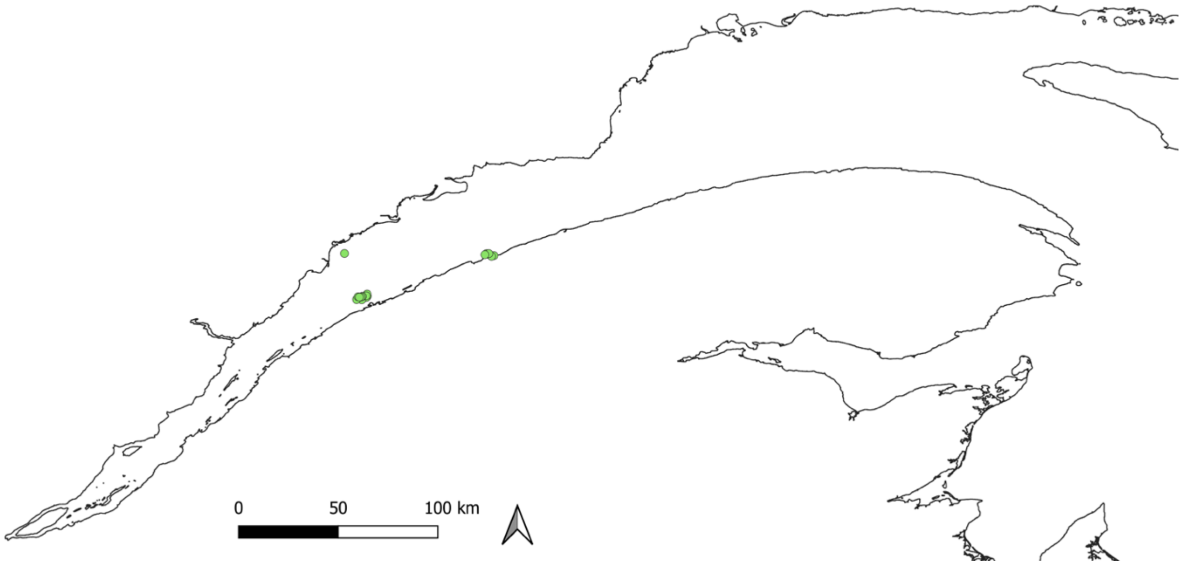


**Figure S2.** Detailed sample location map for Eastern Quebec showing sampling sites per individual (green).

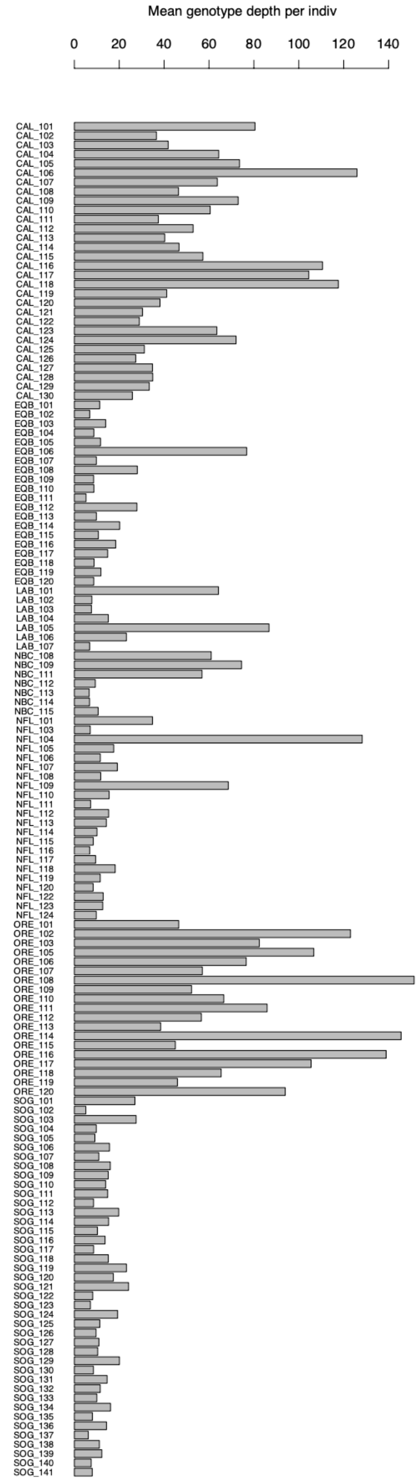


**Figure S3.** Average per locus genotype depth per individual for all individuals in the filtered input VCF from Stacks. Includes 146 individuals genotyped at 10,847 variants from the single SNP per locus dataset.

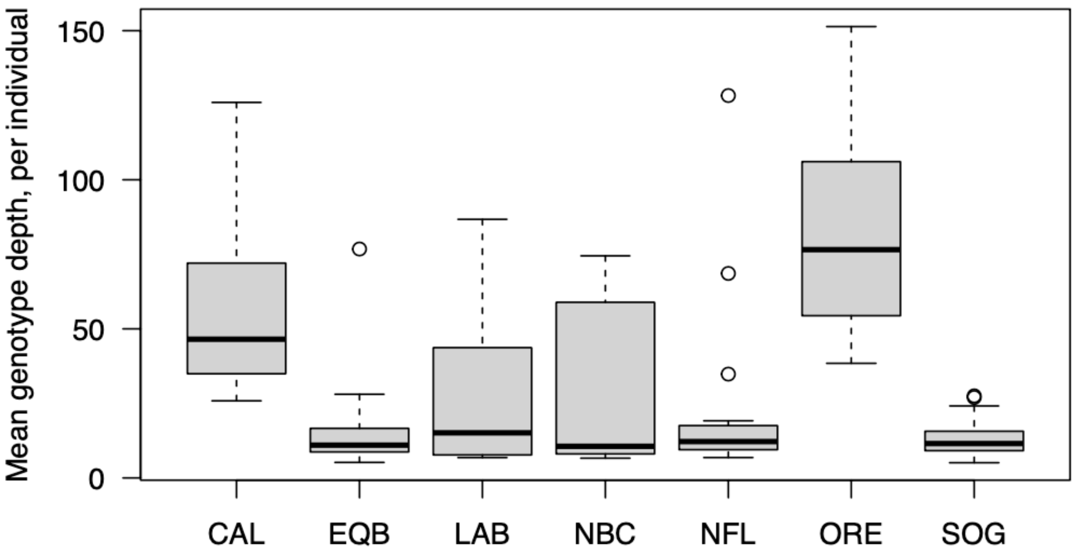


**Figure S4.** Average per locus genotype depth per individual summarized by collection. Abbreviations: CAL = California; EQB = eastern Quebec; LAB = Labrador; NBC = Northern British Columbia (BC); NFL = Newfoundland; ORE = Oregon; SOG = Strait of Georgia, BC.

**(A)**

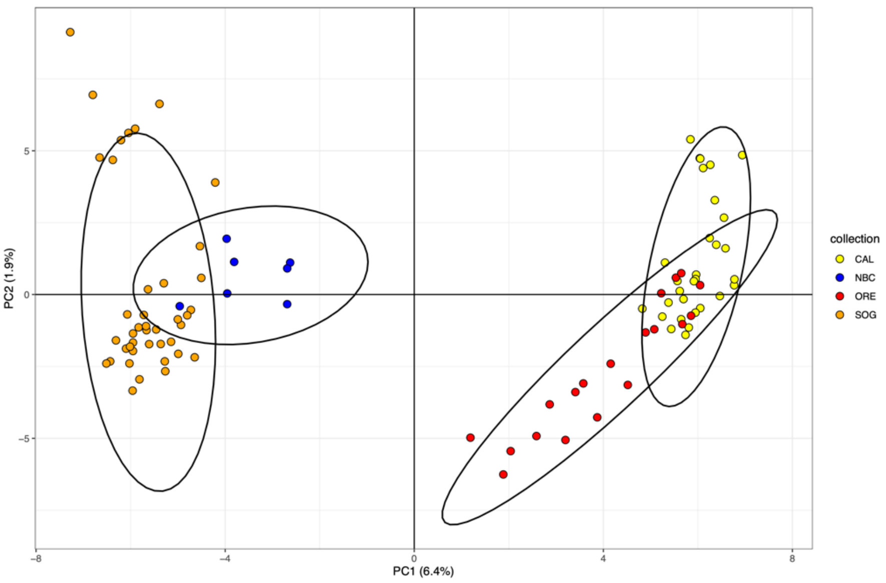


**(B)**

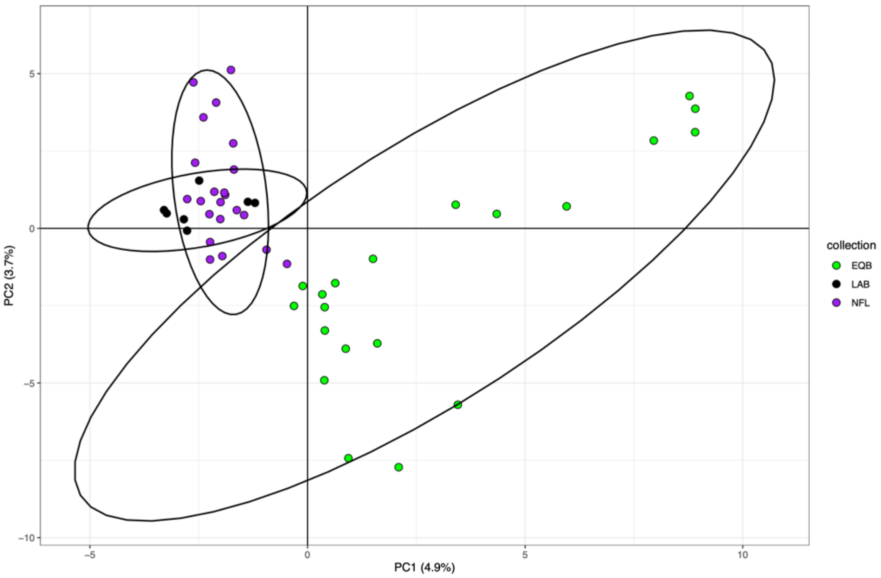


**Figure S5**. Principal components analysis of (A) Pacific collections with 7,707 filtered variants; and (B) Atlantic collections with 3,204 filtered variants, prior to the removal of putative relatives. An outlier cluster is observed in SOG and CAL, which were largely removed by removal of elevated relatedness individuals.

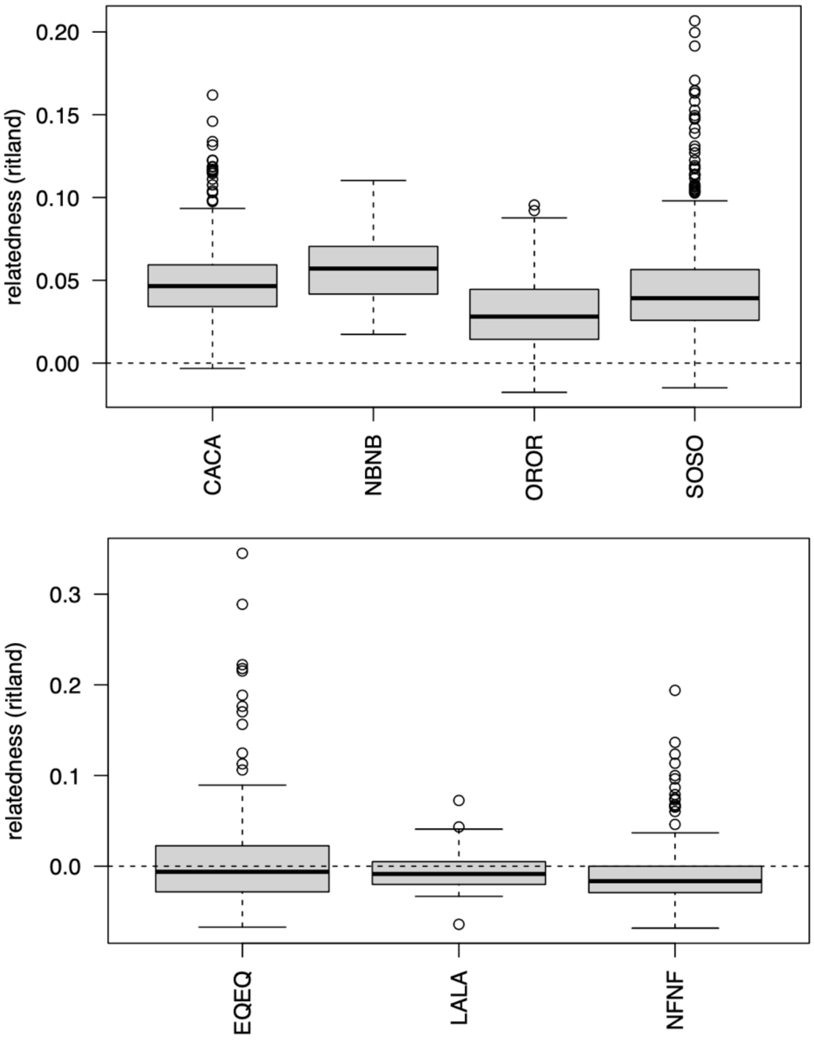


**Figure S6.** Genetic relatedness of within-population pairs in the Pacific (A) and Atlantic (B) datasets calculated within coast-specific datasets. Results are shown prior to the removal of elevated relatedness individuals. High relatedness outlier pairs are indicated as points outside of the 95% confidence limits of the boxplot. Abbreviations for same-on-same comparisons: CACA = California; NBNB = Northern British Columbia (BC); OROR = Oregon; SOSO = Strait of Georgia, BC; EQEQ = eastern Quebec; LALA = Labrador; NFNF = Newfoundland.

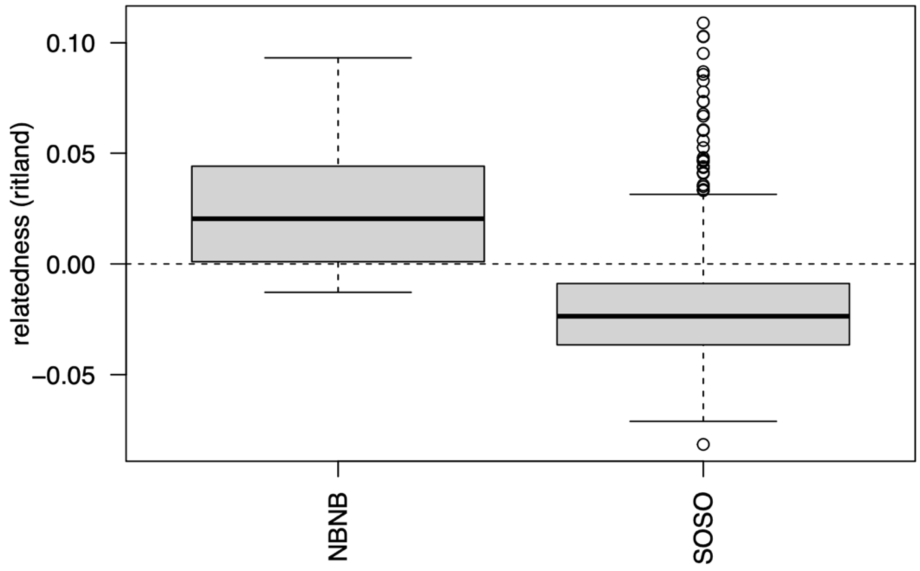


**Figure S7.** Genetic relatedness of within-population pairs when calculated using a BC-only dataset. The value of the Ritland relatedness statistic was reduced relative to that calculated using all Pacific populations simultaneously, but the identity of the outlier pairs remains largely consistent in both analyses. Abbreviations for same-on-same comparisons: NBNB = Northern British Columbia (BC); SOSO = Strait of Georgia, BC.

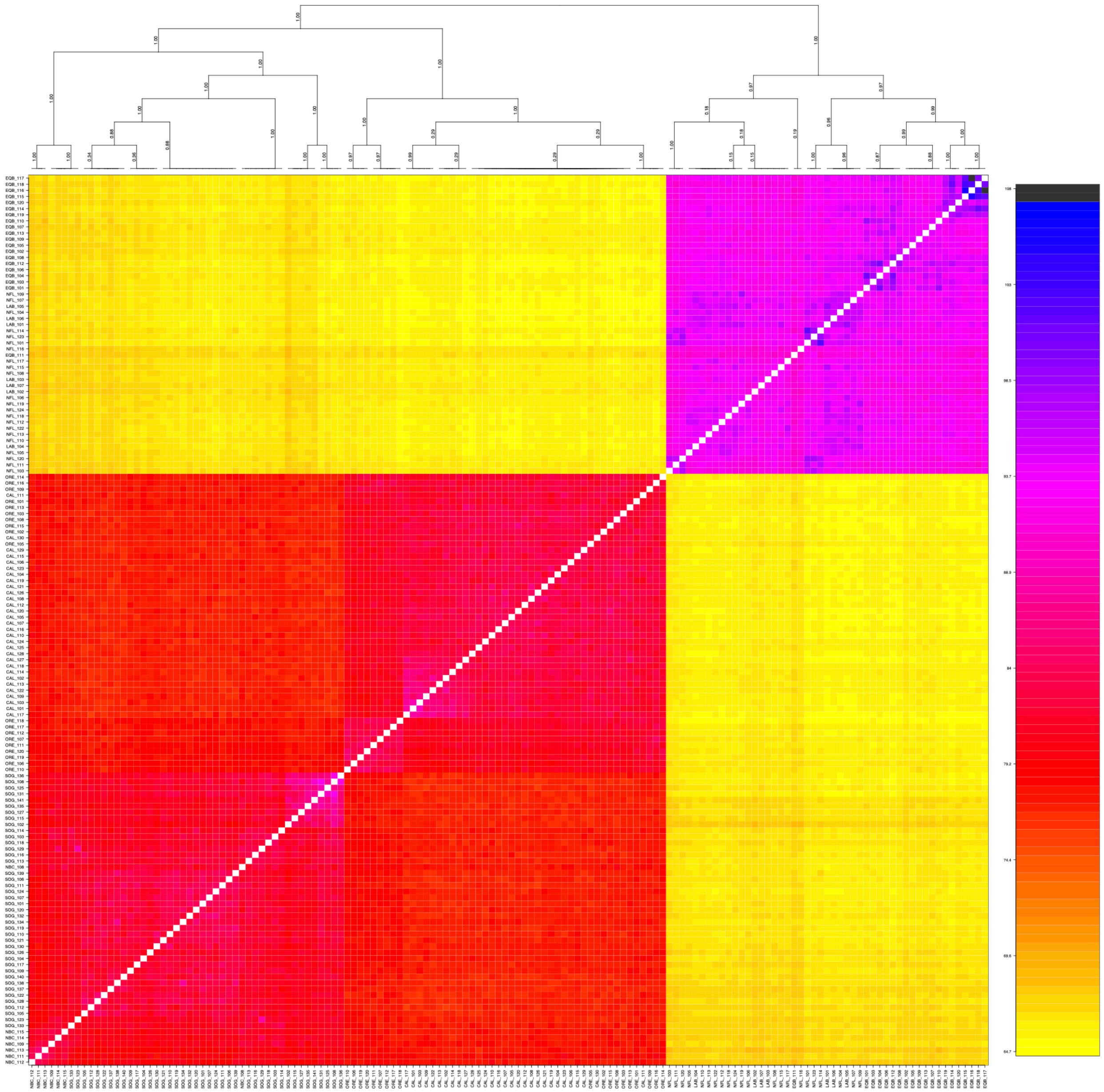

**Figure S8.** Shared microhaplotypes and similarity among individuals clustered into groupings showing the strong dissimilarity between subspecies, and also clusters of individuals with elevated genetic relatedness. Values at tree nodes indicate the proportion of trees that show the clustering pattern represented by the tree.

**(A)**

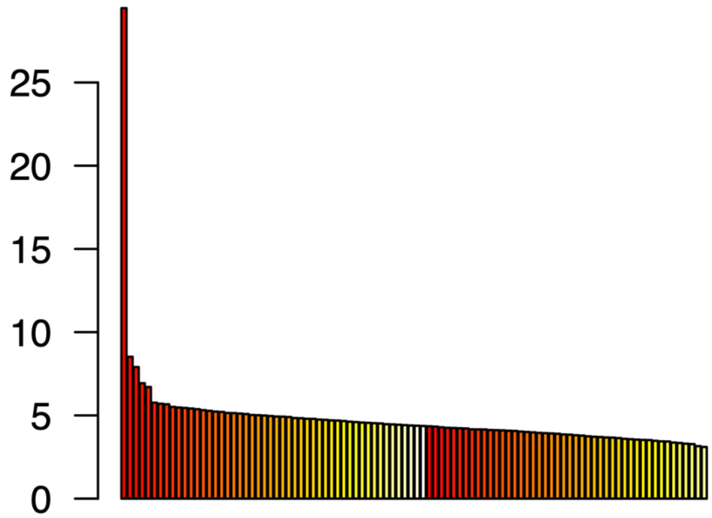


**(B)**

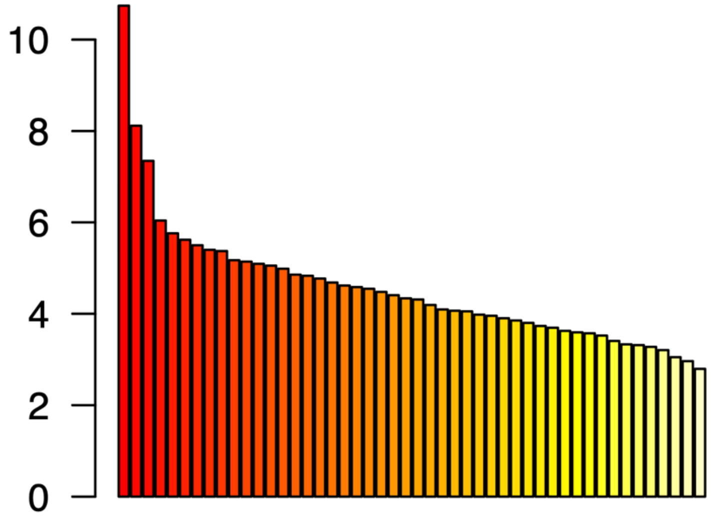


**Figure S9.** Scree plots for the principal components analysis of the coast-specific datasets (with putative elevated related individuals retained) for (A) the Pacific dataset; and (B) the Atlantic dataset.

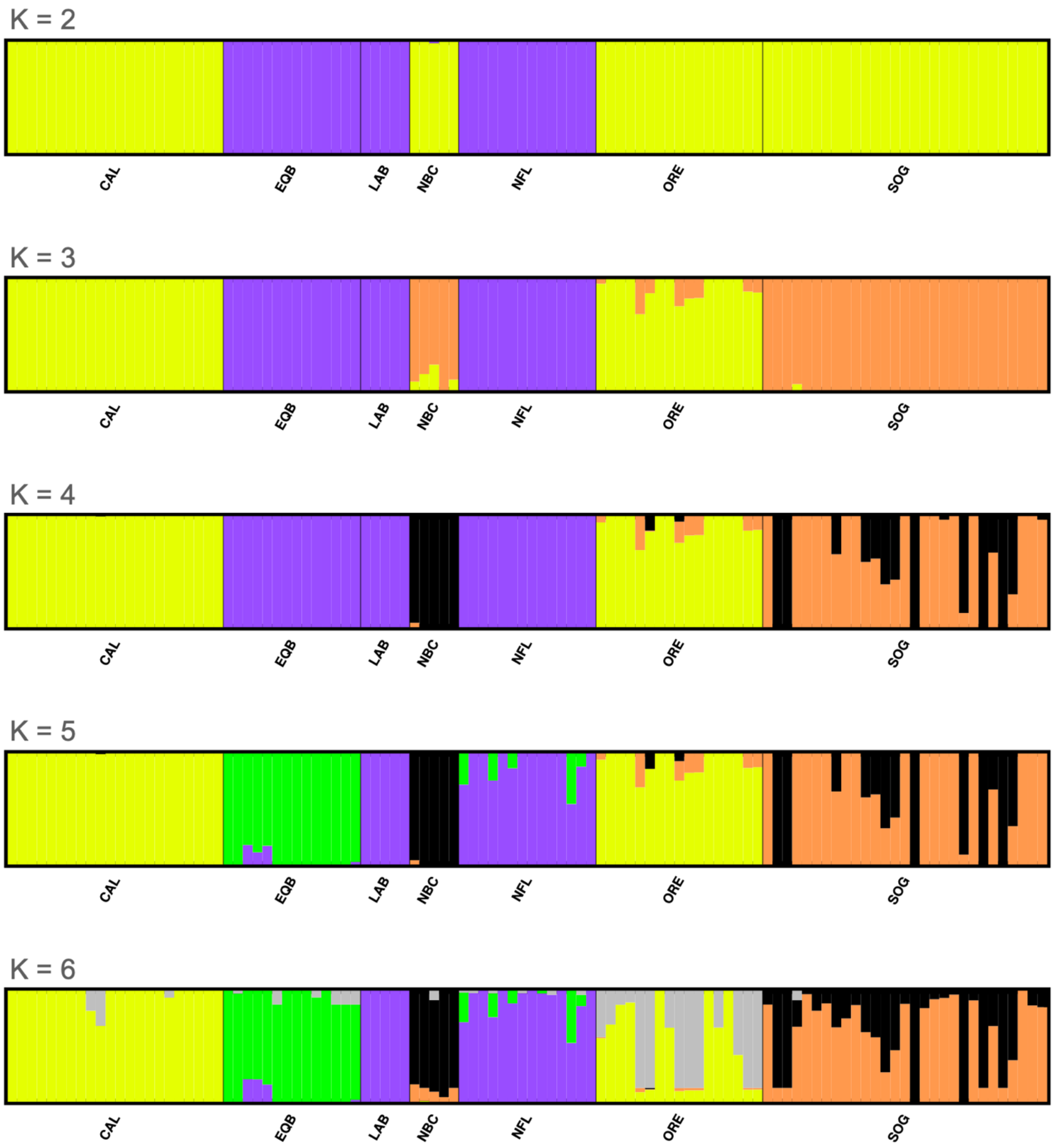


**Figure S10.** Per individual admixture proportions for the major modes for *K*=2 through *K*=6 using both coasts and all samples after putative close relatives were removed.

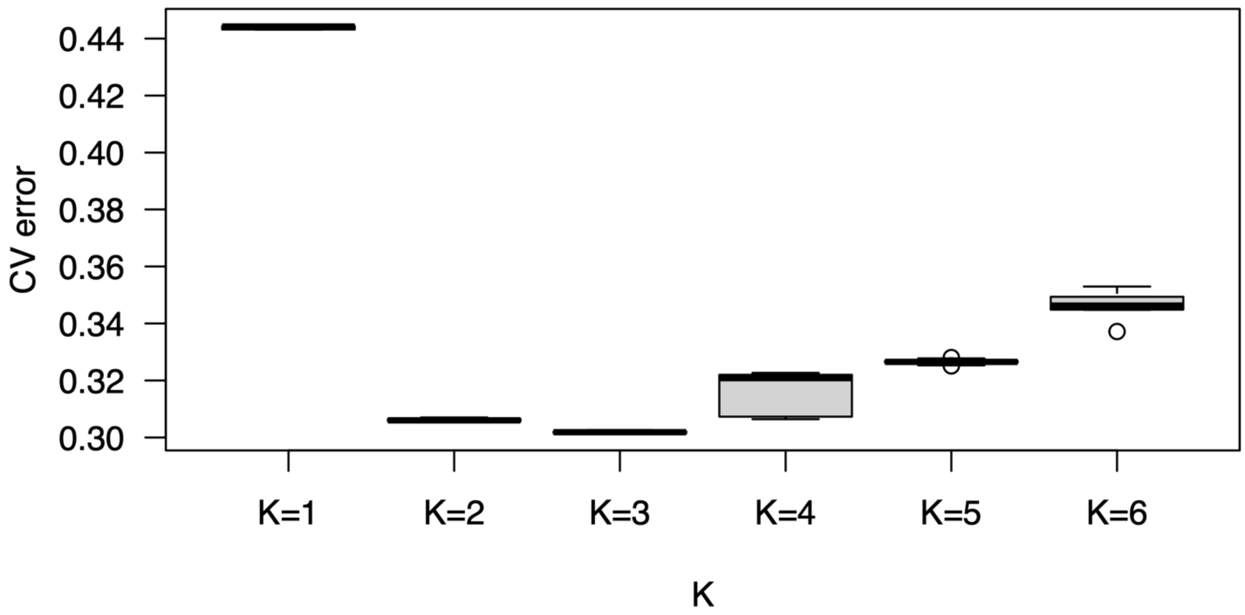


**Figure S11.** Cross validation error for *K*=1 through *K*=6 with ten runs per value of *K* using both coasts and all samples after putative close relatives were removed.

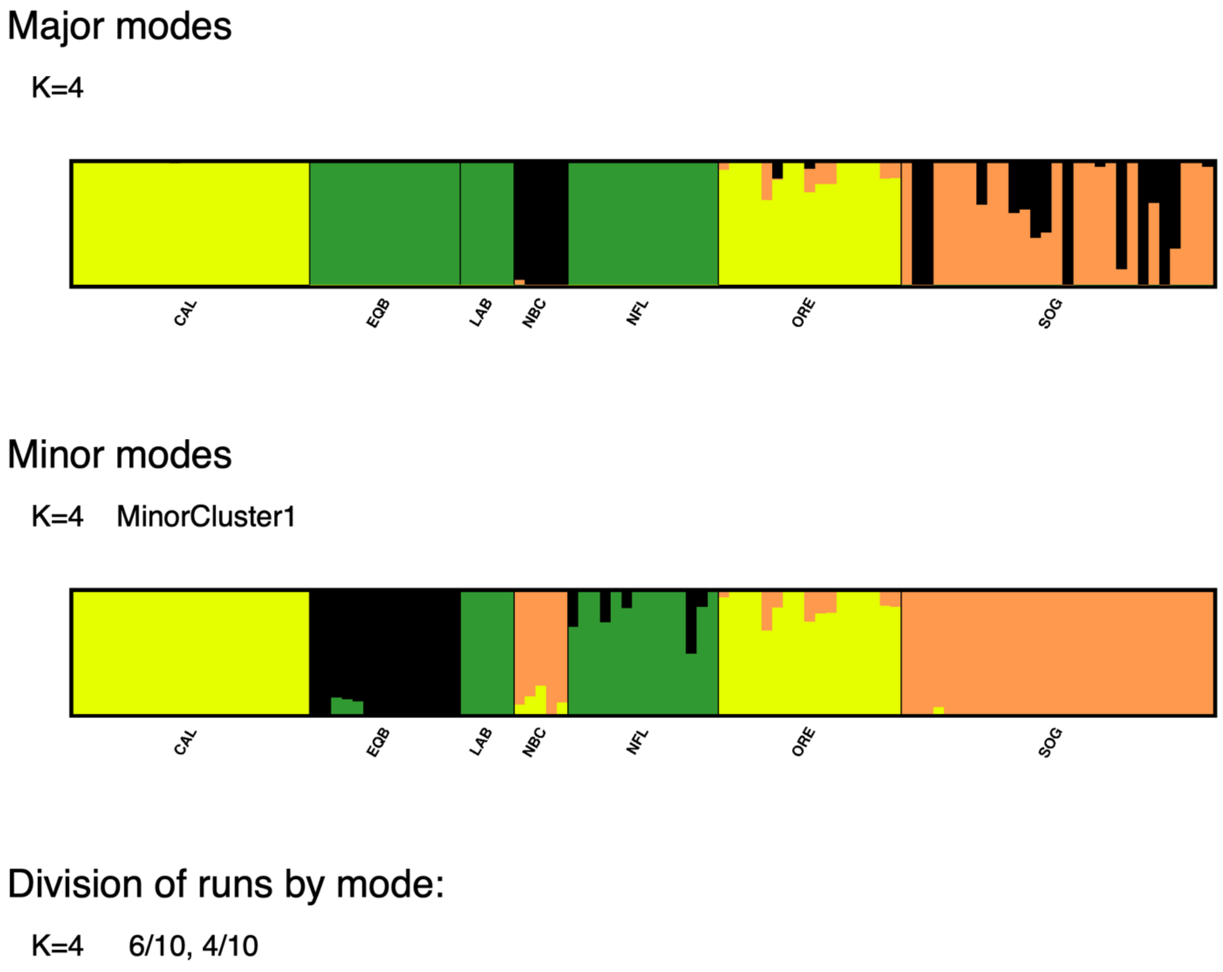


**Figure S12.** Per individual admixture proportions for major and minor modes for *K*=4 using both coasts and all samples after putative close relatives were removed.

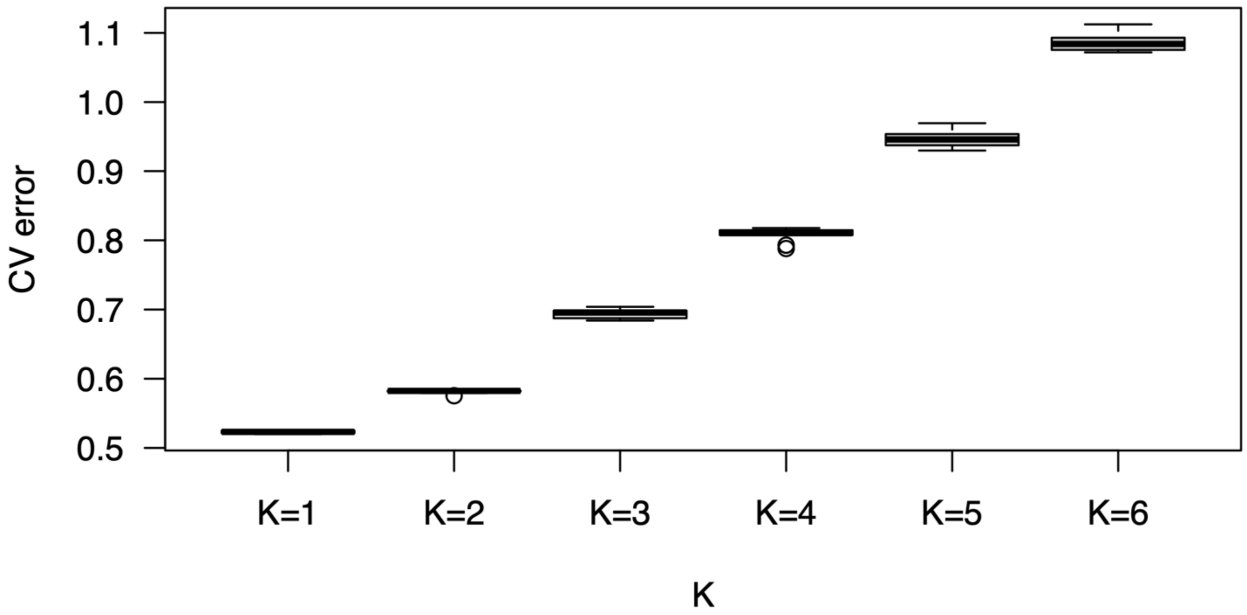


**Figure S13.** Cross validation error for *K*=1 through *K*=6 with ten runs per value of *K* for east coast collections only after putative close relatives were removed.

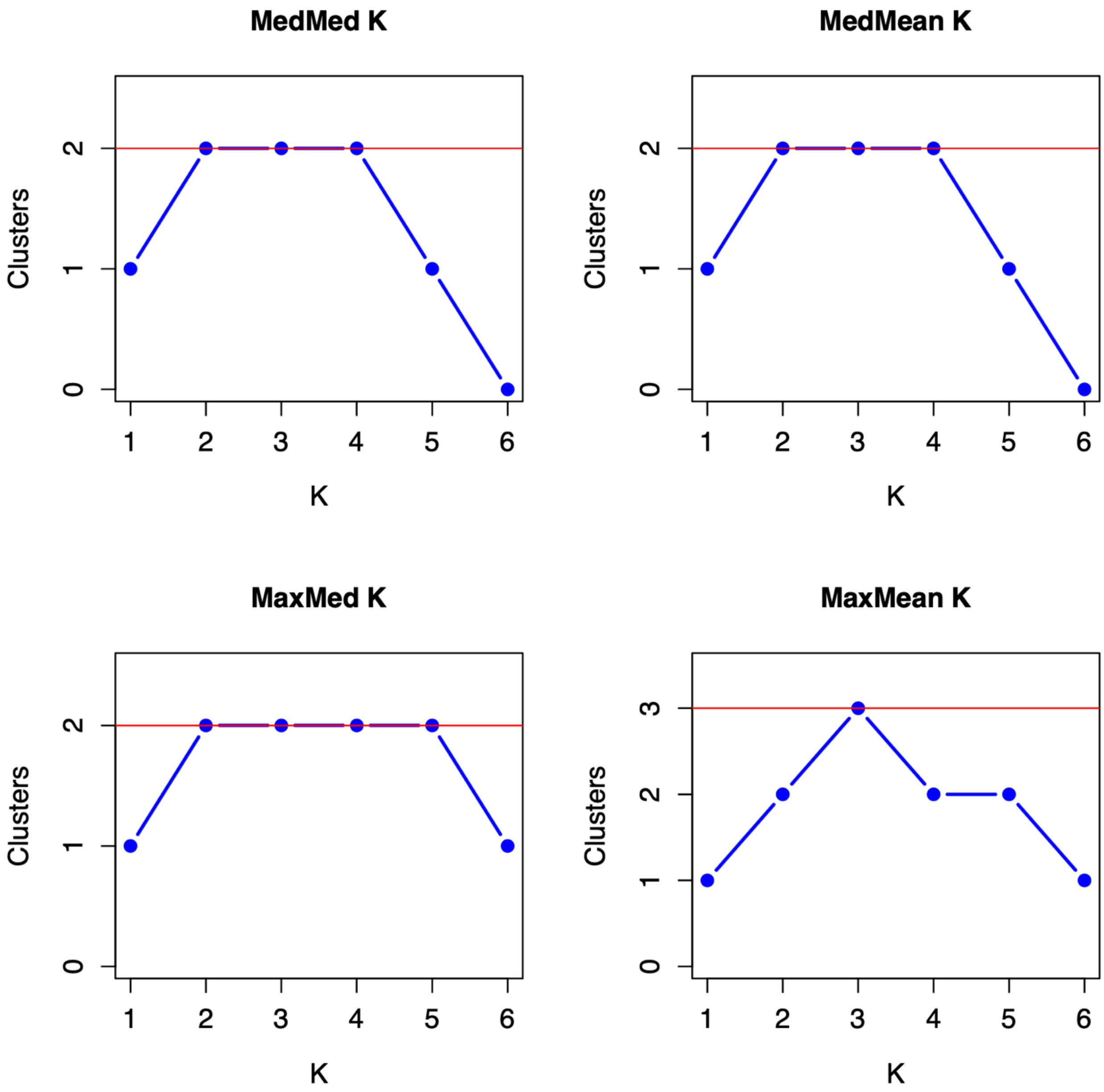


**Figure S14.** Optimal *K* selection metrics for the east coast dataset using StructureSelector.

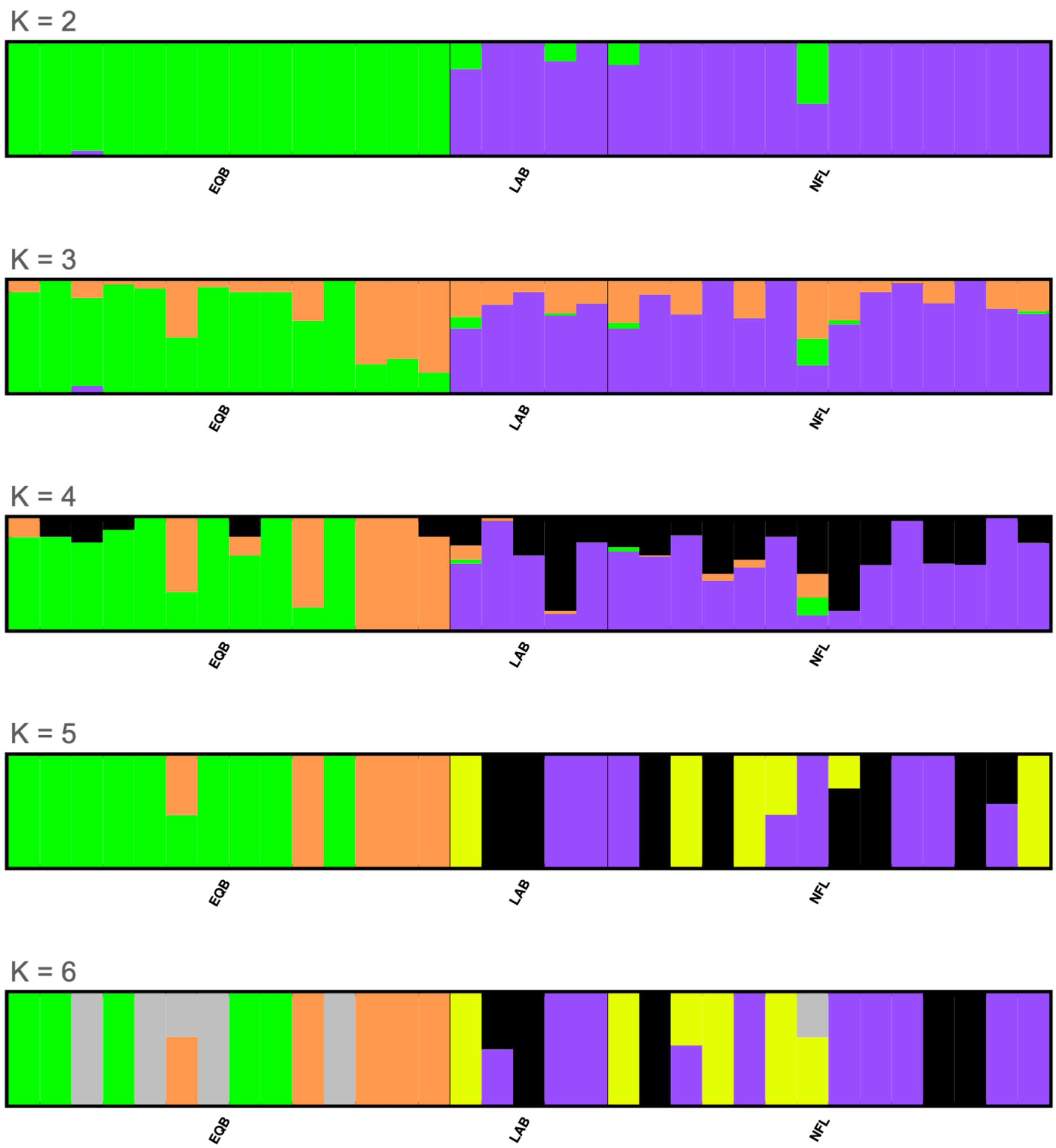


**Figure S15.** Per individual admixture proportions for the major modes for K=1 through K=6 using the east coast collections only coasts and all samples after putative close relatives were removed.

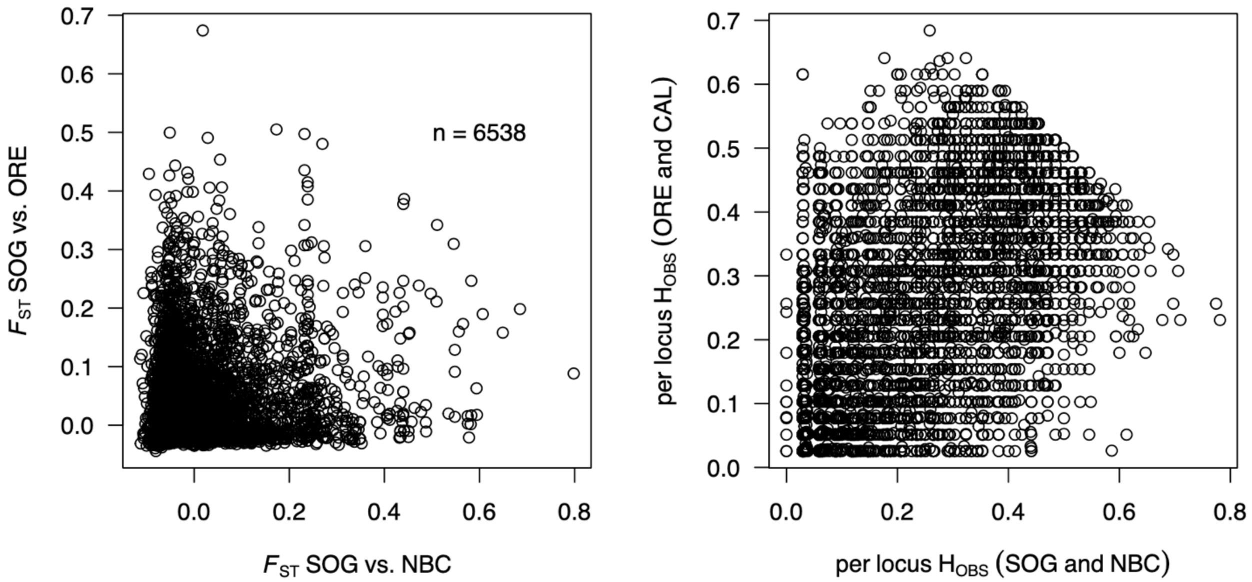


**Figure S16.** (A) Genetic differentiation ($F_{ST}$) per locus for contrasts of SOG vs. ORE, and SOG vs. NBC indicates that some markers have high $F_{ST}$ in both comparisons. Variants included (n = 6,538) had MAF > 0.01 in each data subset. (B) Observed heterozygosity ($H_{OBS}$) per marker was calculated for the USA populations (ORE and CAL), and separately for the BC populations (SOG and NBC), then compared to each other to identify elevated $H_{OBS}$ in both regions. Marker names and associated per locus statistics for each contrast are provided in the Additional Files.

**Table S1.** Collection permit details for all collection regions in the study.

| Collection | Permit Details |
|---|---|
| **NBC** | Samples were collected by DFO under DFO permits, and by the Vancouver Aquarium Marine Mammal Rescue Center under a Statement of Work with DFO. |
| **SOG** | Samples were collected by DFO under DFO permits, and by the Vancouver Aquarium Marine Mammal Rescue Center under a Statement of Work with DFO. |
| **ORE** | Samples were provided by the Oregon Marine Mammal Stranding Network collected under Oregon State University's Stranding Agreement with NOAA Fisheries. |
| **CAL** | Samples were provided by the Marine Mammal Center of Sausalito, California collected under a Stranding Agreement with NOAA Fisheries. |
| **EQB** | Samples were collected by DFO under DFO permits. |
| **NFL** | Samples were collected by DFO under DFO permits, or by hunters licensed under the Fisheries Act. |
| **LAB** | Samples were collected by DFO under DFO permits, or by hunters licensed under the Fisheries Act. |

**Table S2.** All population $F_{ST}$ results after removal of putative relatives. The 95% confidence intervals of $F_{ST}$ are shown with the lower limit in the lower half of the grid and upper limit in the upper half of the grid.

| | CAL | EQB | LAB | NBC | NFL | ORE | SOG |
|---|---|---|---|---|---|---|---|
| **CAL** | - | 0.422 | 0.384 | 0.062 | 0.417 | 0.008 | 0.068 |
| **EQB** | 0.401 | - | 0.030 | 0.481 | 0.020 | 0.427 | 0.410 |
| **LAB** | 0.362 | 0.019 | - | 0.429 | 0.002 | 0.384 | 0.377 |
| **NBC** | 0.053 | 0.457 | 0.405 | - | 0.475 | 0.050 | 0.020 |
| **NFL** | 0.397 | 0.014 | -0.006 | 0.451 | - | 0.422 | 0.406 |
| **ORE** | 0.005 | 0.406 | 0.362 | 0.042 | 0.401 | - | 0.055 |
| **SOG** | 0.062 | 0.389 | 0.355 | 0.014 | 0.385 | 0.050 | - |

**Table S3.** Datasets A (present study) and B (Liu et al. 2022) with associated statistics to evaluate the effect of the Pacific-based reference genome on diversity metrics. $H_{OBS}$ is based on all positions (variants and fixed). Dataset B *de novo* used trimmed length reads where reads were required to be of constant length, and therefore did not use the full dataset.

| Data-set | Genotyping Approach | Pop | Avg. n per locus | Reads (M) per pop | Align (M) per pop | Obs. het ($H_{OBS}$) | Pi | Total (Mbp) | Poly-morphic loci | % Poly |
|---|---|---|---|---|---|---|---|---|---|---|
| A | Reference-based, all pops | NBC | 6.8 | 30.3 | 27.6 | 0.00044 | 0.00044 | 4.646 | 6400 | 0.138 |
| | | SOG | 39.8 | 138.6 | 122.7 | 0.00043 | 0.00044 | | 9018 | 0.194 |
| | | ORE | 18.8 | 104.5 | 99.5 | 0.00046 | 0.00046 | | 8358 | 0.180 |
| | | CAL | 29.7 | 108.7 | 102.8 | 0.00045 | 0.00046 | | 8528 | 0.184 |
| | | EQB | 19.5 | 73.8 | 63.9 | 0.00024 | 0.00024 | | 3875 | 0.083 |
| | | LAB | 6.9 | 29.0 | 25.6 | 0.00025 | 0.00024 | | 3257 | 0.070 |
| | | NFL | 21.4 | 76.8 | 66.5 | 0.00024 | 0.00024 | | 4044 | 0.087 |
| | *de novo*, Pacific only | SOG | 15.0 | 22.5 | *NA* | 0.00039 | 0.0004 | 1.399 | 2434 | 0.174 |
| | | ORE | 16.4 | 22.5 | *NA* | 0.00048 | 0.00048 | | 2692 | 0.192 |
| | *de novo*, Atlantic only | EQB | 15.0 | 22.5 | *NA* | 0.00024 | 0.00021 | 3.273 | 2733 | 0.084 |
| | | NFL | 14.7 | 22.5 | *NA* | 0.00024 | 0.00022 | | 2831 | 0.086 |
| B | Reference-based, all pops | KOD | 13.1 | 10.9 | 10.8 | 0.00034 | 0.00037 | 7.281 | 10760 | 0.148 |
| | | END | 12.6 | 7.2 | 7.1 | 0.00031 | 0.00034 | | 10409 | 0.143 |
| | | BIC | 13.9 | 21.7 | 21.3 | 0.00019 | 0.00019 | | 5573 | 0.077 |
| | | NFL | 13.6 | 18.6 | 18.3 | 0.00019 | 0.00019 | | 5744 | 0.079 |
| | | ORK | 13.8 | 18.2 | 18.2 | 0.00010 | 0.00009 | | 2087 | 0.029 |
| | | WNL | 13.0 | 10.2 | 10.1 | 0.00009 | 0.00009 | | 2772 | 0.038 |
| | *de novo*, Pacific only | KOD | 11.5 | 7.3 | *NA* | 0.00074 | 0.00058 | 0.299 | 662 | 0.221 |
| | | END | 10.7 | 4.7 | *NA* | 0.00074 | 0.00059 | | 644 | 0.215 |
| | *de novo*, west Atlantic only | BIC | 13.1 | 14.7 | *NA* | 0.00041 | 0.00035 | 3.226 | 3832 | 0.119 |
| | | NFL | 11.8 | 12.9 | *NA* | 0.00042 | 0.00037 | | 4215 | 0.131 |
| | *de novo*, east Atlantic only | ORK | 13.0 | 14.0 | *NA* | 0.00037 | 0.00026 | 1.231 | 911 | 0.074 |
| | | WNL | 11.6 | 7.3 | *NA* | 0.00036 | 0.00026 | | 1183 | 0.096 |

**Table S4.** Samples that were removed due to being in a pair over the relatedness cutoff to remove putative close relatives from the analysis.

| Collection | Number removed | Removed individuals due to putative relatedness |
|---|---|---|
| **NBC** | 2 | NBC_112, NBC_113 |
| **SOG** | 12 | SOG_137, SOG_108, SOG_115, SOG_136, SOG_141, SOG_128, SOG_130, SOG_129, SOG_132, SOG_131, SOG_135, SOG_138 |
| **ORE** | 2 | ORE_119, ORE_120 |
| **CAL** | 8 | CAL_103, CAL_113, CAL_114, CAL_117, CAL_122, CAL_126, CAL_121, CAL_123 |
| **EQB** | 6 | EQB_103, EQB_104, EQB_116, EQB_117, EQB_118, EQB_120 |
| **NFL** | 8 | NFL_103, NFL_111, NFL_114, NFL_120, NFL_123, NFL_119, NFL_124, NFL_122 |
| **LAB** | 2 | LAB_105, LAB_103 |
